# How marine currents and environment shape plankton genomic differentiation: a mosaic view from *Tara* Oceans metagenomic data

**DOI:** 10.1101/2021.04.29.441957

**Authors:** Romuald Laso-Jadart, Michael O’Malley, Adam M. Sykulski, Christophe Ambroise, Mohammed-Amin Madoui

## Abstract

Plankton seascape genomics show different trends from large-scale weak differentiation to micro-scale structures. Prior studies underlined the influence of environment and seascape on a few single species differentiation and adaptation. However, these works generally focused on few single species, sparse molecular markers, or local scales. Here, we investigate the genomic differentiation of plankton at macro-scale in a holistic approach using *Tara* Oceans metagenomic data together with a reference-free computational method to reconstruct the *F*_ST_-based genomic differentiation of 113 marine planktonic species using metavariant species (MVS). These MVSs, modelling the species only by their polymorphism, include a wide range of taxonomic groups comprising notably 46 Maxillopoda/Copepoda, 24 Bacteria, 5 Dinoflagellates, 4 Haptophytes, 3 Cnidarians, 3 Mamiellales, 2 Ciliates, 1 Collodaria, 1 Echinoidea, 1 Pelagomonadaceae, 1 Cryptophyta and 1 Virus. The analyses showed that differentiation between populations was significantly lower within basins and higher in bacteria and unicellular eukaryotes compared to zooplantkon. By partitioning the variance of pairwise-*F_ST_*matrices, we found that the main drivers of genomic differentiation were Lagrangian travel time, salinity and temperature. Furthermore, we classified MVSs into parameter-driven groups and showed that taxonomy poorly determines which environmental factor drives genomic differentiation. This holistic approach of plankton genomic differentiation for large geographic scales, a wide range of taxa and different oceanic basins, offers a systematic framework to analyse population genomics of non-model and undocumented marine organisms.

## Introduction

Marine species from epipelagic plankton are drifting organisms abundantly present in every ocean, playing an active role in Earth biogeochemical cycles (1,2) ◻ and form a complex trophic web (3,4) of high taxonomic diversity (5–7) at the basis of fish resources (8,9)◻. Understanding the present connectivity between populations or communities of plankton is thus crucial to apprehend upheavals due to climate change consequences in oceans (10,11)◻.

Due to their potential high dispersal and huge population size, planktonic species have long been thought to be homogenous and highly connected across oceans, but this assumption is challenged by empirical studies since two decades (12)◻. Planktonic species are characterized by theoretical high population effective sizes (13,14)◻, which reduces the power of drift and makes selection and beneficial mutation stronger drivers of their evolution, as exampled in the SAR11 alphaproteobacteria (15)◻, but the balance between neutral evolution and selection is still debated (16,17)◻. Furthermore, evolution in plankton also seems to be strengthened by acclimation through variation of gene expression or changing phenotypes in response to environmental conditions (18–21)◻.

Gene flow and connectivity between planktonic populations can be impacted by three major forces: marine currents, abiotic (i.e physico-chemical parameters) and biotic factors. First, as planktonic species are passively and continuously transported by marine currents, we could expect that isolation-by-distance shapes the genetic structure of populations. Conversely, cosmopolitan, panmictic and/or unstructured species have been reported multiple times in Copepoda (21–24)◻, Collodaria (25)◻ or Cnidaria (26). Other studies show more complex patterns, with genetic structure mainly observed at the level of basins in Copepoda (27)◻, Pteropoda (28)◻, Diatoms (29)◻ and Cnidaria (30) or at mesoscale in Chaetognatha (31)◻, Copepoda (32–34)◻, Dinophyceae (35) or *Macrocystis pyrifera* (36)◻. Thus, due to the complexity of oceanic processes, classical landscape genomics frameworks began to be applied and adapted (37) to better model the dispersion of populations over seascape, or what we would call “isolation-by-currents”. Hence, modelling oceanic circulation at macro- and meso-scale is a prerequisite to capture the water masses connectivity (38)◻. Successful approaches using data derived from larval dispersal models were used in fish and coral (39–41)◻ and the relatively recent use of Lagrangian travel time estimates combined with genetic data showed promising results (34,36) to better explain gene flow◻.

At the same time, changing environmental conditions may lead to selective pressure that counter the effect of dispersion induced by marine currents, leading to a higher differentiation. The best examples are temperature-driven structures from bacteria to cnidaria (15,30)◻ or the effect of salinity in diatoms (42)◻, that can even favours speciation in estuaries (43)◻. Finally, biotic drivers based on competition and co-evolution were also reported to shape evolution (44)◻. However, abiotic and biotic parameters are often linked to oceanic circulation, which leads to technical challenge to disentangle the role and importance of each parameter on populations’ connectivity.

All these above mentioned findings usefully enhanced our understanding of plankton connectivity, like in zooplankton (45)◻, but they focused on documented species with reference sequences, often using few molecular markers such as mitochondrial (COI) or ribosomal genes (16S, 18S, 28S), and/or are restricted to mesoscale sampling. Thus, we need to overcome these case studies by adopting a holistic approach which integrates the analyses of genome-wide markers belonging to species from different levels of the trophic chain, sampled across the world oceans.

Advances in environmental genomics realized by shotgun sequencing offer a new perspective for population genomics of marine plankton species based on metagenomic data. Diversity in ocean microorganisms can now be better understood, thanks to ambitious expeditions (46,47). Particularly, *Tara* Oceans data provide a unique dataset from many locations in all the world oceans, enabling global approaches to investigate plankton (7,48–51), but blind spots in term of taxonomy or function are still an obstacle for further analyses, due to the lack of reference genomes or transcriptomes. The first way to address this issue relies on the use of the metagenome-assembled genomes (MAGs) from metagenomic data that enable to retrieve a large amount of lineages from metagenomic samples, especially for small-sized genomes as found in viruses and prokaryotes (48,52–55). A second way is the single-cell sequencing after flow-cytometric sorting (56) which allows the genome reconstruction of small eukaryotic species. Both ways increase the number of available references. An alternative way is based on a reference-free approach of metagenomic data (57), in order to analyse the population differentiation of numerous unknown species potentially lacking a reference.

Here, we proposed to study plankton connectivity from a holistic point of view, using metagenomic data extracted from samples gathered during *Tara* Oceans expeditions in Mediterranean Sea, Atlantic and Southern Oceans. After extracting polymorphic data and clustering them into metavariant species (MVS) using a reference-free method (57), we coupled environmental parameters and a new modelling of Lagrangian travel times (58) to estimate the relative contribution of environment and marine currents on the population differentiation of these MVSs.

## Material and Methods

### Extracting metavariants from *Tara* Oceans metagenomic data

Metavariants are nucleotidic variants detected directly from metagenomic data using the reference-free variant caller *DiscoSNP++* (59) with parameters −k 51 -b 1(60) (Arif et al. 2018)◻. We used a set of 23×10^6^ metavariants produced in a previous study (60)◻. These metavariants were detected in 35 *Tara* Oceans sampling sites corresponding to four distinct size fractions (0.8-5 μm, 5-20 μm, 20-180 μm and 180-2000 μm) from the water surface layer, for a total of 114 samples (Figure 1A). For further analyses, *Tara* stations were separated into four groups corresponding to the basins they belong to: the Mediterranean Sea (MED; TARA_7 to TARA_30), Northern Atlantic Ocean (NAO; TARA_4, TARA_142 to TARA_152), Southern Atlantic Ocean (SAO; TARA_66 to TARA_81), and Southern Ocean (SO; TARA_82 to TARA_85). Full protocols for sampling, extractions and sequencing are detailed in previous studies (61,62).

**Figure 1:**
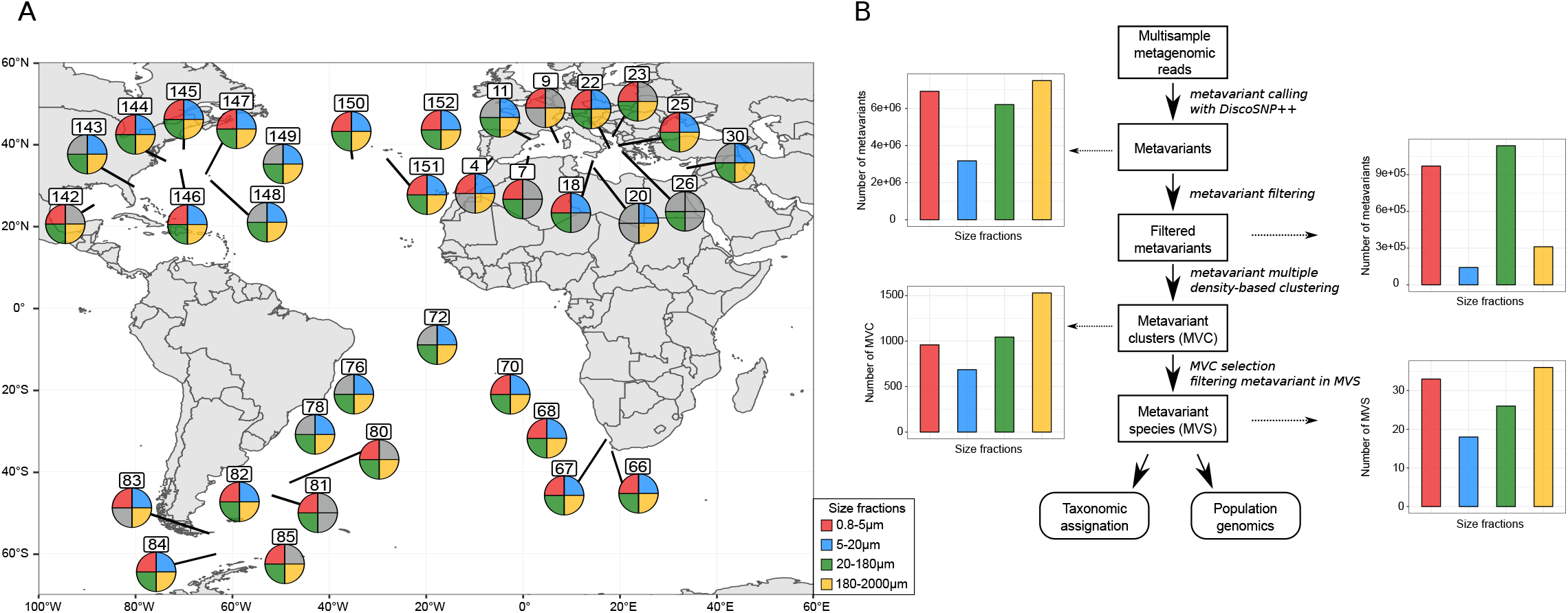
Construction of metavariant species from metagenomic dataset of *Tara* Oceans. A) Worldmap showing the locations of the 35 *Tara* Oceans stations used in the study. Each circle is divided in four, depending on the detection of an MVS. In grey, no MVSs were retrieved. B) Pipeline of MVS construction, with additional statistics by size fraction. From top to bottom: number of metavariants before and after filtering, number of metavariant clusters (MVC) detected and number of metavariant species (MVS) finally selected.

### Construction of metavariant species

To identify sets of loci belonging to unique species, we used metaVaR version v0.2 (57). This method enables the clustering by species of metavariants previously called from metagenomic raw data. Each cluster is constituted of genomic variants of a single species and the final clusters are called metavariant species (MVSs).

The metavariants of the four size fractions were filtered using metaVarFilter.pl with parameters −a 5 −b 5000 −c 4. This process discarded low covered loci, repeated regions that present very high coverage and loci with non-null coverage in less than four samples.

The second step of the metaVaR process clusters the metavariants. MetaVaR uses multiple density-based clustering (dbscan, (63,64)◻), a total of 187 couples of parameters epsilon and minimum points (ε, MinPts) were tested, with epsilon ε = {4,5,6,7,8,9,10,12,15,18,20} and MinPts ={1,2,3,4,5,6,7,8,9,10,20,50,100,200,300,400,500}. This clustering phase constitutes a set of clusters called metavariant clusters (MVC) for each couple (Supplementary Figure S1). Then a maximum weighted independent sets (MWIS) algorithm was used on the resulting set of MVCs to select the best non-overlapping clusters, i.e. clusters sharing no metavariants. For the dataset corresponding to the size fraction 20-180μm, 220 MVCs containing more than 90% of the metavariants were discarded to decrease the memory use during the MWIS computation. For each selected MWIS, only loci with a depth of coverage higher than 8x were kept. Finally, only MVSs with at least 100 variants, and for which at least three samples presented a median depth of coverage > 8x were retained, leading to a final set of 113 MVSs. As a result, metaVaR provides a frequency matrix and a coverage matrix across each biallelic locus in each population for each MVS that will be used further for population genomic analyses.

### Taxonomic assignation of MVSs

To provide a taxonomic assignation of each MVS, three different assignations were performed, using different sources of information (Supplementary Figure S2).

First, for each size fraction, the sequences supporting the metavariants were mapped on downloaded NCBI non-redundant database (10/23/2019) with diamond v0.9.24.125 (65)◻, using blastx and parameter −k 10, and the results were filtered based on the E-value (<10^−5^). Then, for each variant, the taxonomic ID and bitscore of each match were kept. A fuzzy Lowest Common Ancestor (LCA) method (66) was used to assign a taxonomy to each sequence, using bitscore as a weight with −r 0.67 −ftdp options. The highest phylogenetic ranks were retained as the best assignation for each sequence. This constituted a first taxonomic assignation of the metavariant sequences. In parallel, the sequences were mapped on MATOU, a unigen catalog based on *Tara* Oceans metatranscriptomic data (50)◻, and on the MMETSP transcriptomic database (67)◻. This constituted three different taxonomic assignations of the variant sequences.

Then, for each MVS, the unfiltered variant sequences from the corresponding MVC were used to maximize information. The three mentioned taxonomic assignations were crossed with the MVC sequences and the sequences assigned to the same clade were summed and used as a basis for a manual taxonomic assignation of the MVS. Each MVS was thus assigned to the most probable taxonomic clade. MVSs were then regrouped into 24 taxonomic groups that were clustered into six reliable wider groups: Virus, Bacteria, Unicellular Eukaryotes, Animals, Copepods, and Poor classification (Figure 2B). This offered three levels of assignation, from the most precise to the widest (Supplementary Table S1).

**Figure 2:**
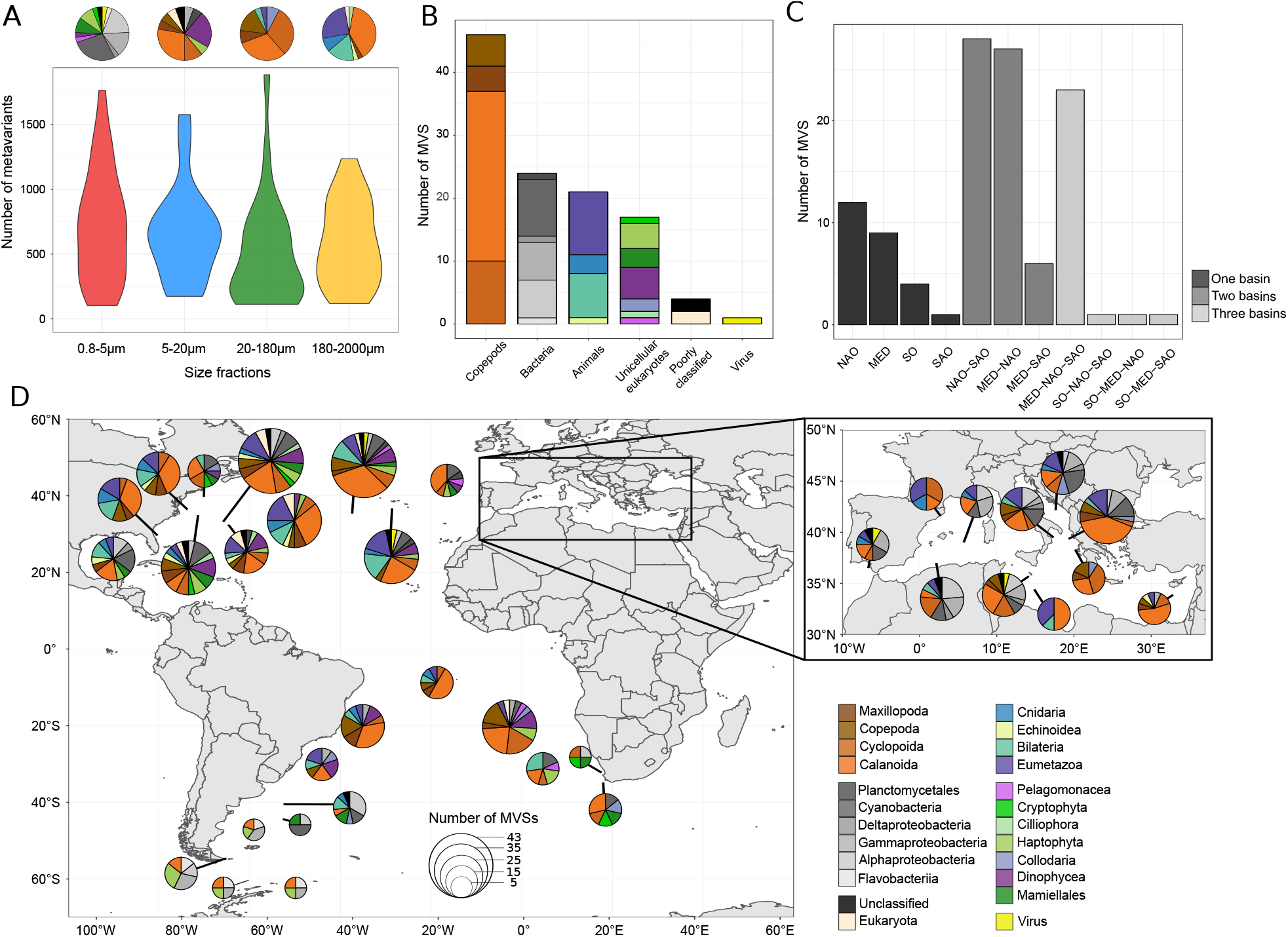
Description of the set of MVSs. A) Distribution of the number of metavariants for each size fraction. On the top, pie charts representing the taxonomic composition of each size fractions. B) Number of MVSs assigned to the six wider taxonomic groups. C) Number of MVSs according to the basins they were detected in: Northern Atlantic Ocean (NAO), SAO (Southern Atlantic Ocean), SO (Southern Ocean) and MED (Mediterranean Sea). D) World map showing the number of MVSs of each taxonomic group for each *Tara* station. The size of the circles corresponds to the amount of MVSs detected in each station. Colors of taxonomic groups are indicated on the bottom right of the panel.

### Population genomics analysis

To investigate genomic differentiation at different scales, the *F*_ST_ metrics was used throughout this study and computed for each variant of an MVS as follows, 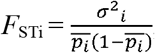, with 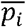 and 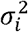 being respectively the mean and variance of allele frequency across the considered populations *i* (68)◻. Two types of computations were launched, in each MVS. A first global *F*_ST_ was calculated using the total set of populations, allowing the analysis of the global *F*_ST_ distribution. Then, a pairwise-*F*_ST_ was calculated between the populations, and median pairwise-*F*_ST_ was retained as a measure of genomic differentiation between the populations of the MVS.

For the whole set of MVSs, each pairwise-*F*_ST_ comparison was extracted from the metaVaR outputs. These pairwise-*F*_ST_ were compared in three different statistical frameworks, by grouping them based on the following factors: the basins where the two populations are located, the taxonomic assignation of the MVS and the size fraction of the MVS. For each comparison, a Kruskal-Wallis test was used to assess the significance of the variation of the median pairwise-*F*_ST_ among groups. When the test was significant (p-value <0.05), multiple comparison Wilcoxon tests were performed between groups.

### Connection within and between basins

To estimate the connection between and within basins, we regrouped *Tara* stations based on their locations (i.e. MED, NAO, SAO and SO), and computed the mean *F_ST_* between and within basins. As an example, if we compared MED to SO, we extracted, from the median pairwise-*F_ST_* matrices of all MVSs, all the median pairwise-*F_ST_* between a MED station and an SO station were compared, and kept the mean of this distribution as an estimate of differentiation.

### Lagrangian travel time estimation

To estimate Lagrangian transport, we used a method based on drifter data (58)◻. The method is used to compute the travel time of the most likely path between *Tara* stations, back and forth. We used the public database of the Global Drifter Program (GDP), managed by the National Oceanographic and Atmospheric Administration (NOAA) (https://www.aoml.noaa.gov/phod/gdp/) containing information from drifters ranging from February 15, 1979 to September 31, 2019. We extracted the data for both drogued and undrogued drifters (i.e. drifters that lost their sock) to maximize the information used by the method. No drifters have ever been observed to get out of the Mediterranean Sea through the Strait of Gibraltar, therefore to avoid missing data, we arbitrarily added 100 years to the travel times of pathways out of the Mediterranean Sea over the Strait of Gibraltar and added 1 year to the pathways going into the Mediterranean Sea, based on previous models on surface water (69,70)◻. We used 450 rotations within the method to reduce the reliance of travel times on the grid system used. Two travel times are obtained by the method for each pair of stations: back and forth, resulting in an asymmetric travel time matrix between all possible station pairings. For our analyses, we retained only the minimum of these two travel times in the matrix, as this then accounts for the direction of currents between stations.

### Environmental data

Environmental variables corresponding to the 35 selected *Tara* stations were extracted from the World Ocean Atlas public database (https://www.nodc.noaa.gov/OC5/woa13/woa13data.html), for the period 2006-2013 on 1°×1° grid, covering the dates of *Tara* Oceans expeditions. The following parameters were retrieved: temperature (°C), salinity (unitless), silicate (μmol.L^−1^), phosphate (μmol.L^−1^) and nitrate (μmol.L^−1^).

### Variation partitioning of the genomic differentiation of MVSs

To estimate the relative contribution of environmental parameters and Lagrangian travel time in the variance of each MVS genomic differentiation, a linear mix model (LMM) was applied with R package MM4LMM (71)◻. The model applied was the following; *Y_FST_* = μ + *Zu* + *ε*, where *Y_FST_* is the vector of observations of *F*_ST_ values with a mean *μ, Z* is a known matrix of parameters relating the observations *Y_FST_* to *u*, a vector of independent random effects of zero mean and *ε* is a vector of random errors of 0 means and covariance matrix proportional to the identity (white noise).

For each pairwise-*F*_ST_ matrix, the corresponding matrix of minimum Lagrangian travel time was retrieved. Temperature, salinity, silicate, phosphate and nitrate measures were extracted for all the stations where the MVS is present, and a Euclidean distance was computed between the stations for each of these parameters. The LMM was then applied on pairwise-*F*_ST_ values using the five environmental distances and Lagrangian travel time after scaling, adding a variance of 1 for each explicative variable. To note, we considered the parameters as independent variables. As a result, an estimate of the contribution of each parameter to the total variance of pairwise-*F*_ST_ is obtained. In addition, a fixed effect and a proportion of variance unexplained (corresponding to the noise) is retrieved.

In order to investigate the structure of the MVSs relative to their *F*_ST_ variance decomposition, two principal component analyses (PCA) were then performed. A first one was done on the variance explained by the six variables and the unexplained part of the variance over the 113 MVSs. From this PCA, the unexplained variance of *F*_ST_ (Supplementary Figure S4) was high in most of MVSs, strongly contributing to the first component (37% explained variance). For clarity, a second PCA was conducted by removing the unexplained part of the variance. For both PCAs, correlation of the variables with the components and the contribution (i.e. the ratio of the cos^2^ of each variable on the total cos^2^ of the components) of the variables to the components were extracted. PCAs were performed using FactoMineR v2.3 R package (72,73)◻.

### Clustering MVSs into specific parameters-driven differentiation groups

The variance explained by each factor was used to represent the MVSs with dimensional reduction through t-distributed Stochastic Neighbor Embedding (t-SNE), using Rtsne R package (74) with a perplexity of 5 and 5,000 iterations and we extracted the MVS coordinates. Then, a k-means clustering (K = 8) was performed to identify MVSs with common patterns of explained variance. To identify which set of parameters drives the differentiation of a cluster, we compared the distributions of the explained variance of each parameter within the cluster using a Kruskal-Wallis and a Wilcoxon paired tests (p-value < 0.05).

## Results

### Taxonomy and biogeography of MVSs

We used 23×10^6^ metavariants generated from 114 metagenomics data of 35 *Tara* samples with *DiscoSNP++* in a previous study (60) as input for metaVaR and we constructed a total of 113 MVS out of 4,220 MVCs (Figure 1B, Supplementary Table S1), containing altogether 68,575 metavariants (0.3% of the total, Figure 1B). The taxonomic assignation of the MVS showed a wide range of lineages spanning all the plankton trophic levels, with a predominance of Maxillopoda/Copepoda (46), Bacteria (24) and Eumetazoa (21, comprising three Cnidaria and one Echinodea) (Figure 2B). In Bacteria, we found 9 Cyanobacteria, with 8 MVSs linked to *Synechococcus* and one to *Prochlorococcus*. Other notable eukaryotic species belonged to Dinophycea (5), Haptophyta (4), Mamiellales (3), Collodaria (2), Ciliophora (2), Cryptophyta (1) and Pelagomonadacea (1). Only four MVSs presented a poor assignation (unclassified or Eukaryotes) and one MVS was a virus. In Mamiellales, two MVSs were identified as *Bathyccocusprasinos* and are related to previously observed results from *Tara* Oceans (Supplementary Table S2). The size of MVSs ranged from 114 to 1,767 variants and was unrelated to the size fraction (Figure 1A, Kruskal-Wallis p-value > 0.05). As expected, bacteria dominate smaller size fractions, and Eumetazoa (Cnidaria, Bilateria, Copepods) are found in higher size fractions.

A vast majority of MVSs (95, 84%) were present in four to six stations, with a maximum of eight stations for an MVS (Supplementary Figure S5). The number of MVSs per stations showed an important variation (Figure 2D), from four to 43 MVSs (TARA_67/81/84/85 and TARA_150 respectively). Notably, stations from Southern Ocean (TARA_82 to 85) contained few MVSs compared to the others (from 4 to 7 MVSs), with four MVSs (Gammaproteobacteria, Haptophyta, Flavobacteriia and Calanoida) being solely present in Southern Ocean (SO). Finally, 36 MVSs were present in only one basin, while a majority of MVSs (80) were present in Northern Atlantic Ocean (NAO) and in one other basin (Figure 2C).

### Global view of MVSs genomic differentiation

Pairwise-*F*_ST_ was used to estimate the population differentiation among the MVSs. First, we saw that differentiation between populations was significantly more important among basins than within basins (Figure 3A), for each size fraction separately or together. When we compared the basins (Figure 3B), NAO presented weak differentiation with MED and SAO (0.118 and 0.143 respectively). SAO and MED presented a relatively higher differentiation between them (0.222). Finally, this analysis underlined the important global differentiation of the SO from other basins (0.201-0.555), but also a high differentiation within the SO (0.397).

**Figure 3:**
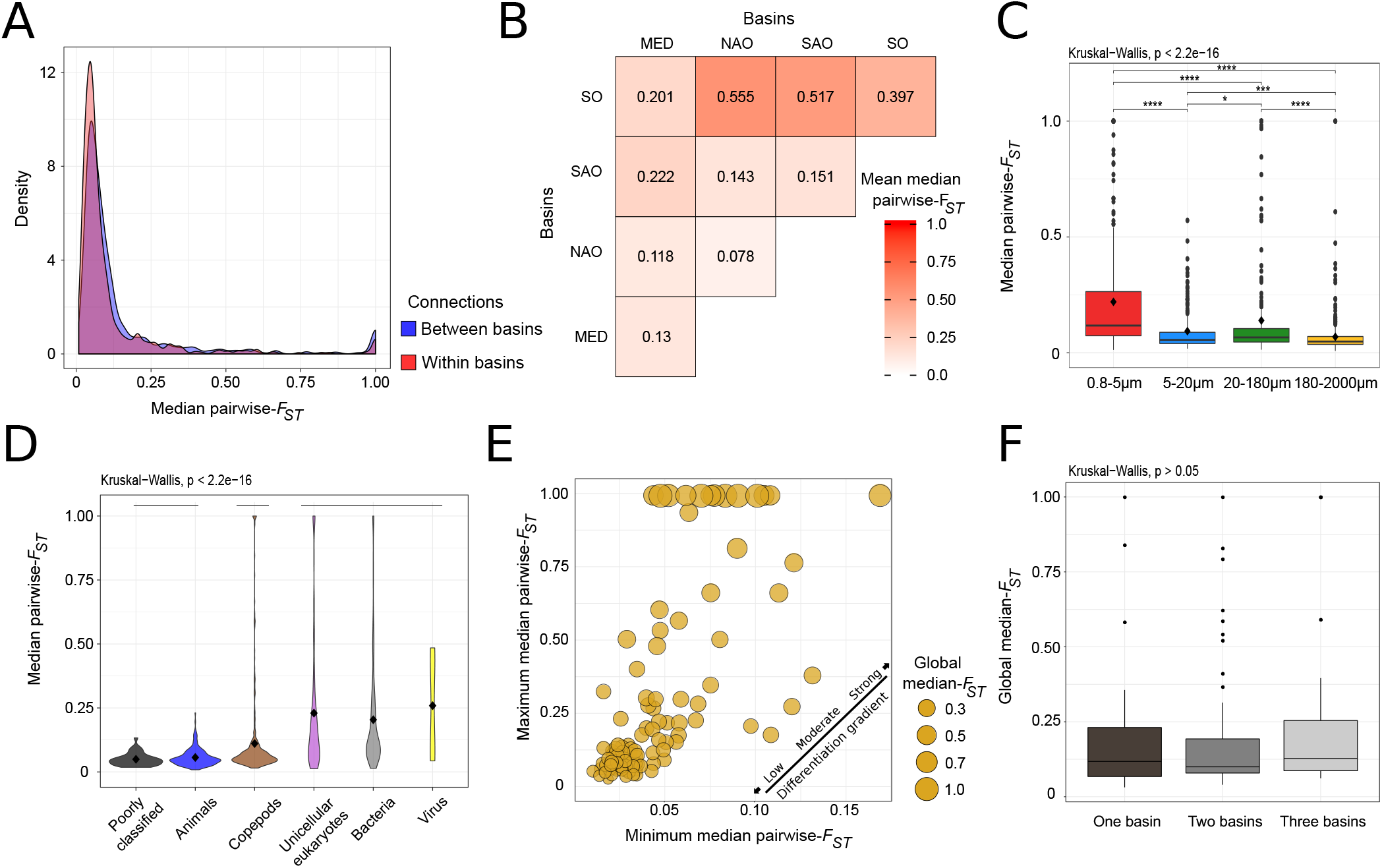
Global view of genomic differentiation. A) Distributions of the 113 MVSs’ pairwise-*F_ST_* matrices. In red, pairwise-*F_ST_* of populations belonging to the same basin; in blue to different basins. B) Pairwise-*F_ST_* matrix between basins. The values represent the mean of all the median-*F_ST_* between stations regrouped according to the basin they belonged to. C) Distributions of the MVSs’ median pairwise-*F_ST_*, according to their size fractions. Black diamonds correspond to the mean of the distributions. The bars on the top correspond to the comparisons done by pairwise Wilcoxon tests (p-values: * <0.05, **<0.01, ***<0.001, ****<0.0001) D) Distributions of the MVSs’ median pairwise-*F_ST_*, according to their taxonomic group. Black diamonds correspond to the mean of the distributions. Each bar corresponds to taxonomic groups displaying no significant differences. E) Scatter plot, each dot is an MVS. The size of each dot reflects the global median-*F_ST_* of the MVS’ *F_ST_* distribution (i.e., *F_ST_* computed over all the populations of an MVS). F) Global median *F_ST_* compared to the number of basins MVSs were detected. Each dot is an MVS.

Secondly, population differentiation was significantly different between size fractions (Kruskal-Wallis, p-value < 0.05), being higher in 0.8-5μm and lower in 180-2000μm (Figure 3C). Population differentiation between the six larger taxonomic groups (see Methods) was related to the body size of the lineages, with a differentiation being relatively lower in copepods and other animals than in unicellular eukaryotes, bacteria and virus (Figure 3D).

We observed a large spectrum of population genomic differentiation patterns among MVSs (Figure 3E), with maximum median pairwise-*F*_ST_ between 0.03 and 1. Extreme cases were observed, for 13 MVSs presenting one or more populations with a median pairwise-*F*_ST_ of 1, and a global *F*_ST_ distribution strongly shifted to 1, as exampled by the Collodaria (MVS 15_200_2, Supplementary Figure S6). We then saw that the number of basins where MVSs were spotted was not significantly linked to their global *F_ST_*(Kruskal-Wallis p-value > 0.05, Figure 3F).

### Computing Lagrangian estimates of marine travel times

Based on recorded drifter motion throughout the ocean, we computed Lagrangian travel time estimates between the 35 *Tara* stations, and observed three clear patterns, distinguishing the MED, NAO and SAO/SO (Figure 4A, Supplementary Figure S7). These results also showed interesting cases illustrated by the following four examples: (i) the relative proximity from TARA_66 to 76 (SAO) and to other NAO stations, (ii) the link from SO stations to TARA_66 and 70, despite a large geographic distance, (iii) the isolation from TARA_145 to the rest of NAO stations, (iv) a separation from TARA_7/9/11 to the rest of MED stations.

### Estimating the relative role of environment and marine currents

To estimate the relative role of environmental factors and marine currents in the genomic differentiation of plankton, we first extracted the data from World Ocean Atlas (Figure 4B) for temperature, salinity, nitrate, silicate and phosphate. Then, we modelled pairwise-*F*_ST_ of each MVS as the variable depending on the five environmental and Lagrangian times variables using a linear mixed model (LMM). The fixed part of the explained variance was low for each MVS, ranging from 0 to 14% (Supplementary Table S1), and was not further analysed. Among all tested environmental variables, Lagrangian travel time, temperature and salinity were the major contributors to the genomic differentiation (Figure 5A), highly correlated to the three first components (67% explained variance). The variance contribution of nitrate, silicate and phosphate respectively followed on the last three components.

**Figure 4:**
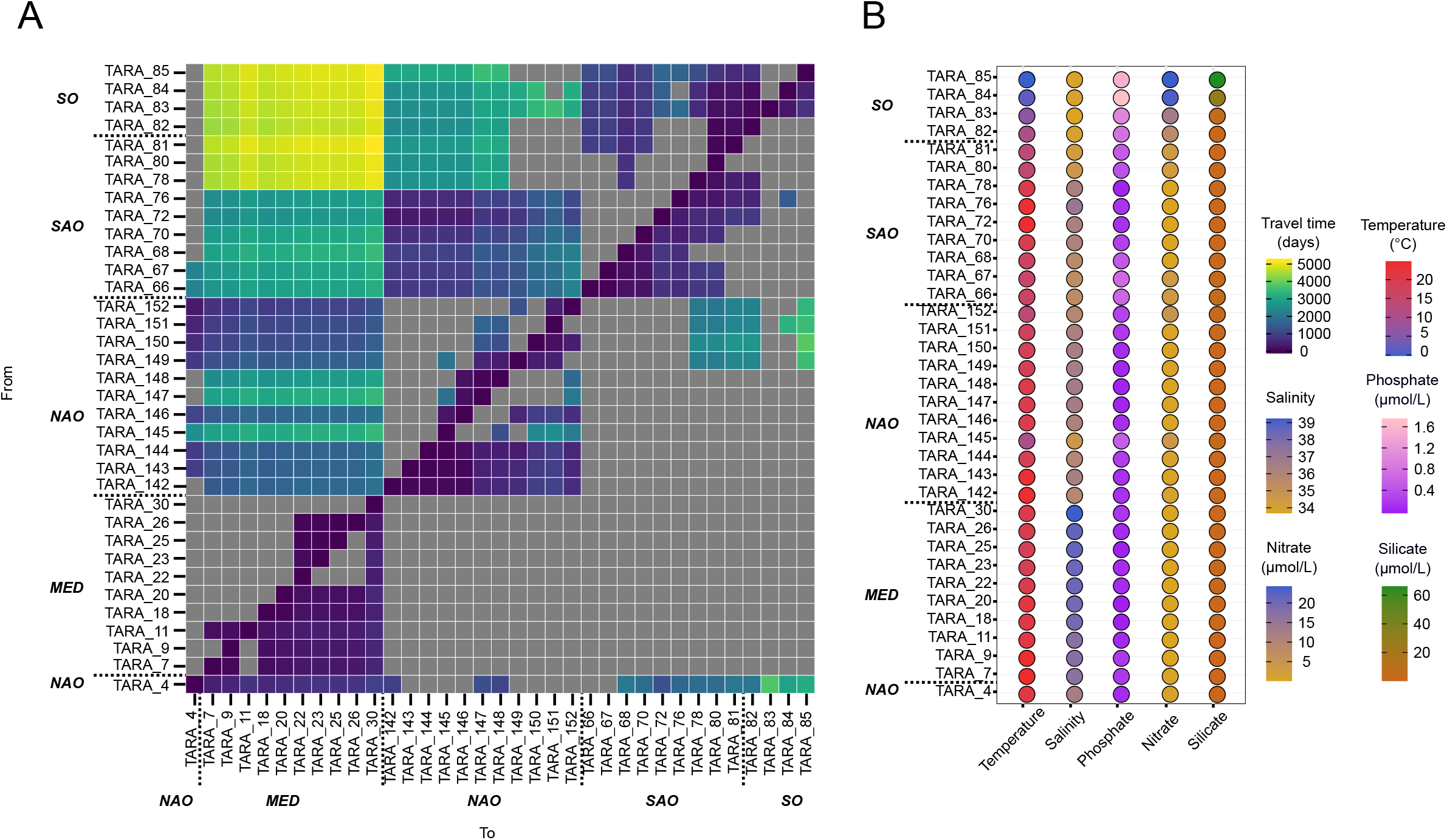
Lagrangian travel times and environmental parameters. A) Minimum times retained for analyses. In grey, asymmetric times that were not the minimum, thus the matrix accounts for the “direction” of currents between stations. B) Measures of temperature, salinity, nitrate, phosphate and silicate extracted from World Ocean Atlas (WOA) for the 35 *Tara* stations. On the right, color scales for each parameter. For the worldmap of *Tara* stations, see supplementary Figure S3.

**Figure 5:**
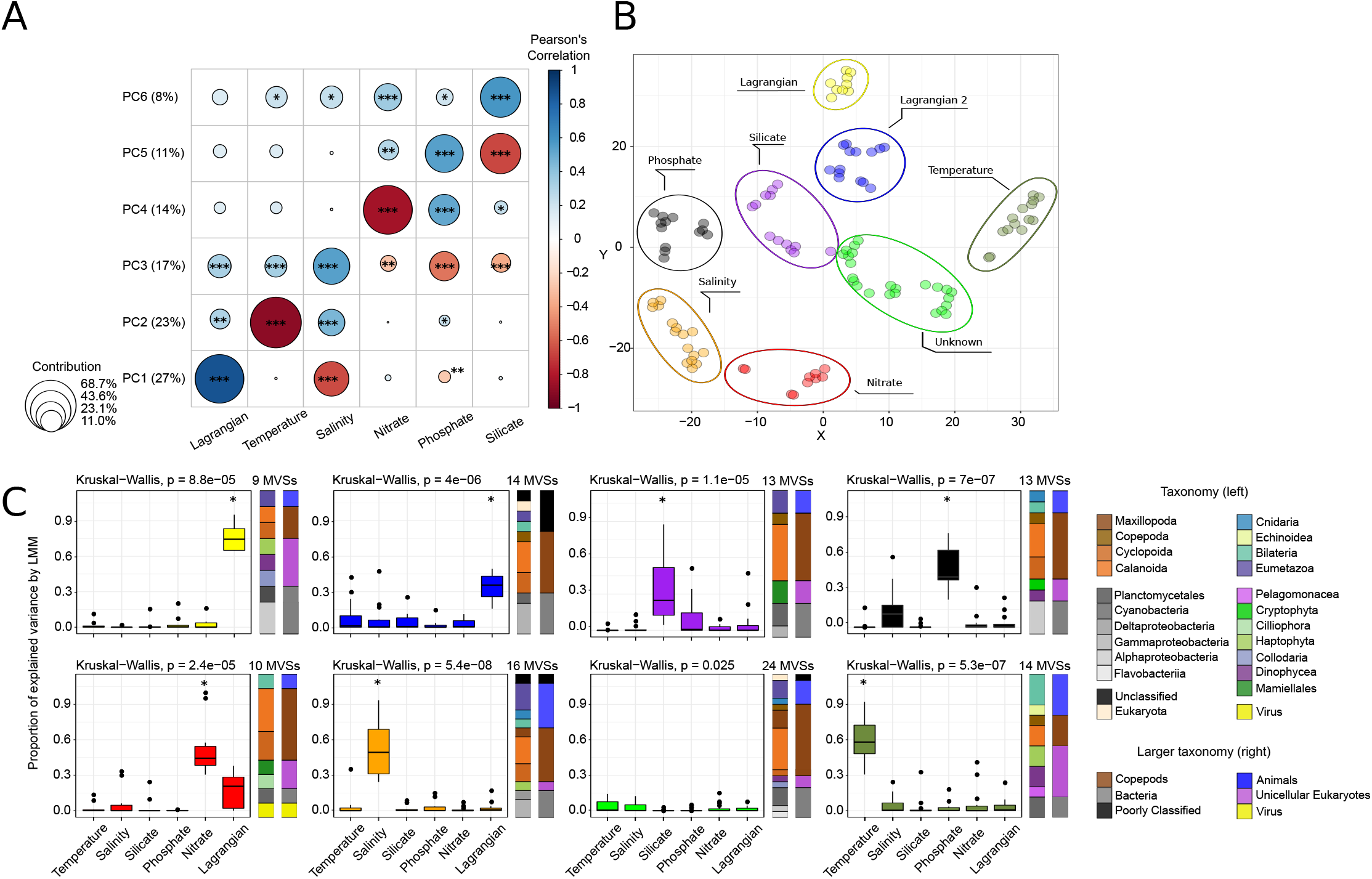
Variation partitioning of genomic differentiation. A) PCA performed on the proportion of variation explained by each parameter over the 113 MVSs. The colour corresponds to the Pearson’s correlation between coordinates of MVSs for a component and the variation explained by the parameters (p-values: * <0.05, **<0.01, ***<0.001, ****<0.0001). The size of the circles represents the relative contribution (i.e. the ratio of the variable cos^2^ on the total cos^2^ of the component) of each variable to each component. B) t-SNE and kmeans (K=8) clustering. Each dot represents an MVS. Each colour corresponds to a defined cluster obtained by kmeans. The names of the clusters are linked to the following figure C) Distributions of variation explained by each factor by cluster, and the taxonomic composition of each cluster. The boxplots colours are the same as the previous figure. The asterisk * on the top of boxplots corresponds to parameters that significantly contributes the most to the genomic differentiation of the MVSs included in the cluster, according to a pairwise Wilcoxon test (p-value < 0.05).

MVSs were then clustered into eight groups by k-means, based on their t-SNE coordinates (Figure 5B). Then, we identified the most important variables over the MVSs of each cluster (Figure 5C), to characterize the clusters. Two clusters were linked to Lagrangian travel times, labelled as “Lagrangian” (14 MVSs) and “Lagrangian 2” (13), the latter exhibiting a lower explained variance by Lagrangian. The largest cluster contained 24 MVSs but was not linked to any parameter. The others are linked to a single environmental parameter: salinity (16 MVSs), temperature (14), silicate (13), phosphate (13) and nitrate(10).

More precisely, the clusters “Lagrangian”, “Temperature” and “Salinity” presented clear differences between their respective drivers compared to the other parameters (Figure 5C). The clusters “Phosphate” and “Silicate” showed a wider distribution of their respective driver among the MVSs they contained, with respectively salinity and phosphate sharing high proportion of explained variance. The “Nitrate” cluster also regrouped MVSs for which a non-negligible part of variance was explained by Lagrangian travel time.

Each cluster showed MVS assigned to almost all taxonomic groups and presented no particular visual enrichment (Figure 5C). This absence of enrichment is clearer in copepods, which constitute the majority of MVSs (Fisher’s Exact Test p-value = 0.348).

Among the nine MVSs belonging to the “Lagrangian” cluster, we observed five MVSs present in Mediterranean Sea and Southern Atlantic and one in Northern and Southern Atlantic. Interestingly, two MVSs were restrained to a single basin, respectively Southern Ocean and Northern Atlantic. Notably, the latter, Planctomycetales 9_200_1, shows a differentiation linked to local marine barriers, with TARA_148 being isolated from the others, TARA_150 and 151 being closely related, and TARA_152 connected to the others, but with slightly higher values (Figure 6A).

**Figure 6:**
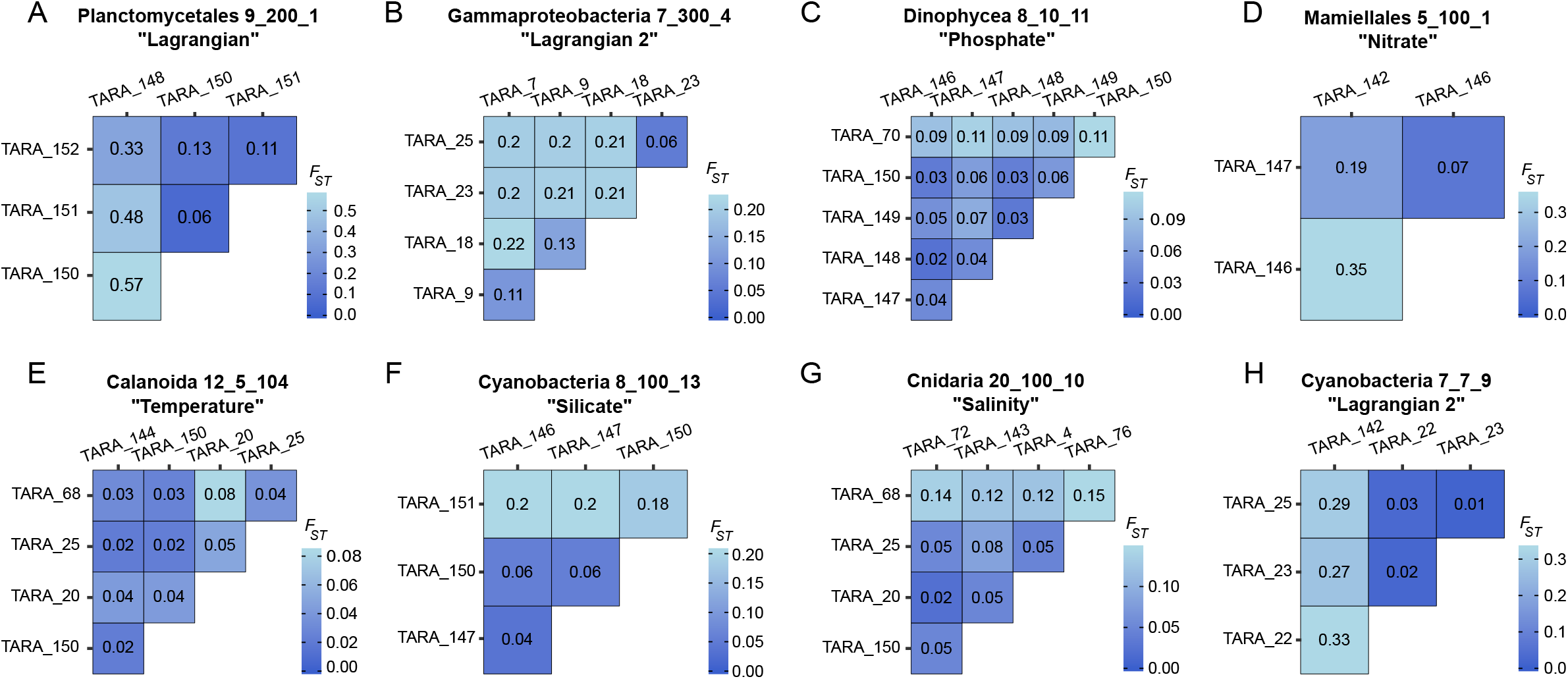
Examples of genomic differentiation. A) to H) Pairwise-*F_ST_*matrices of MVSs mentioned in the respective titles. For each title are mentioned: the taxonomic assignation, the name, and the cluster to which the MVS belongs.

Another example one of within–basin differentiation concerns the Mediterranean gammaproteobacteria 7_300_4 from the “Lagrangian 2” cluster, for which the differentiation clearly shows a pattern linked to marine currents (Figure 6B), with a clear separation between TARA_7, 9 and TARA_23, 25, and TARA_18 being genetically closer to TARA_9, this is explained by Lagrangian estimates together with a small contribution of salinity.

Some MVSs displayed a clear link between their differentiation and one environmental parameter. For example, in the “Phosphate” cluster, we found a Dinophyceae MVS (8_10_11), that displayed a clear unimodal *F_ST_* distribution and no structure between NAO and SAO (Figure 6C). For this Dinophyceae, the population of TARA_70 seemed more isolated to the other NAO populations and TARA_70 is characterized by a higher phosphate concentration (0.264 μmol.L^−1^ against 0.031-0.106 μmol.L^−1^).

Inside the “Nitrate” cluster, there is an example of one Mamiellale MVS (5_100_1) for which populations from TARA_146 and TARA_147 were highly connected, and TARA_142 was more connected to TARA_146 than TARA_147. This reflects the differences in nitrate between these locations (Figure 6D).

In the “Temperature” cluster, the cosmopolitan Calanoida MVS 12_5_104, detected in the MED, NAO and SAO (Figure 6E), presented a relatively higher genetic distance between populations from TARA_20 and 68 (*F_ST_* = 0.08). This genetic pattern was linked to a higher difference in temperature of 5.2°C with respectively 21.9°C and 16.7°C.

In the “Silicate” cluster, we have an illustration of a differentiation along a gradient of silicate, in the cyanobacteria 8_100_13, showing a high isolation of the TARA_151 population compared to populations from TARA_146, 147 and 150 (Figure 6F). The genetic isolation of TARA_151 was linked to a higher concentration in silicate in Northern East Atlantic.

We also found MVSs belonging to a cluster but showing another parameter that also explained a great proportion of the genomic differentiation. As an example, we found the Cnidaria 20_100_10 from the “Salinity” cluster, for which temperature was also an important explaining factor (Figure 6G). Also, the Cyanobacteria 7_7_9 from “Lagrangian 2” cluster presented a clear differentiation between MED and NAO (Figure 6H), which was explained by both Lagrangian travel times and salinity, the Mediterranean Sea presenting higher salinity than NAO.

### Focus on Antarctic genomic differentiation of plankton

From the analysis of global *F_ST_*, it seemed SO presented a pattern of relative isolation from the other basins (Figure 3B). Indeed, we observed that the same four MVSs (Gammaproteobacteria 12_100_16, Flavobacteriia 7_100_6, Haptophyta 4_50_2 and Calanoida 5_20_1) can be found in stations TARA_82, 83, 84 and 85 from the SO. Furthermore, using Lagrangian trajectories (Supplementary Figure S8), the two main currents of the area were spotted: the Malvinas Current and The Antarctic Circumpolar Current (ACC) (Figure 7A). These MVSs presented among the highest global median *F_ST_* (0.35 to 0.84, see Supplementary Figure S6), revealed a very high differentiation between their populations (Figure 7B), and all belonged to different clusters (“Salinity”, “Unknown”, “Lagrangian” and “Nitrate” respectively). Particularly, the Haptophyta MVS presents a differentiation linked to both the ACC and the Malvinas Current.

**Figure 7:**
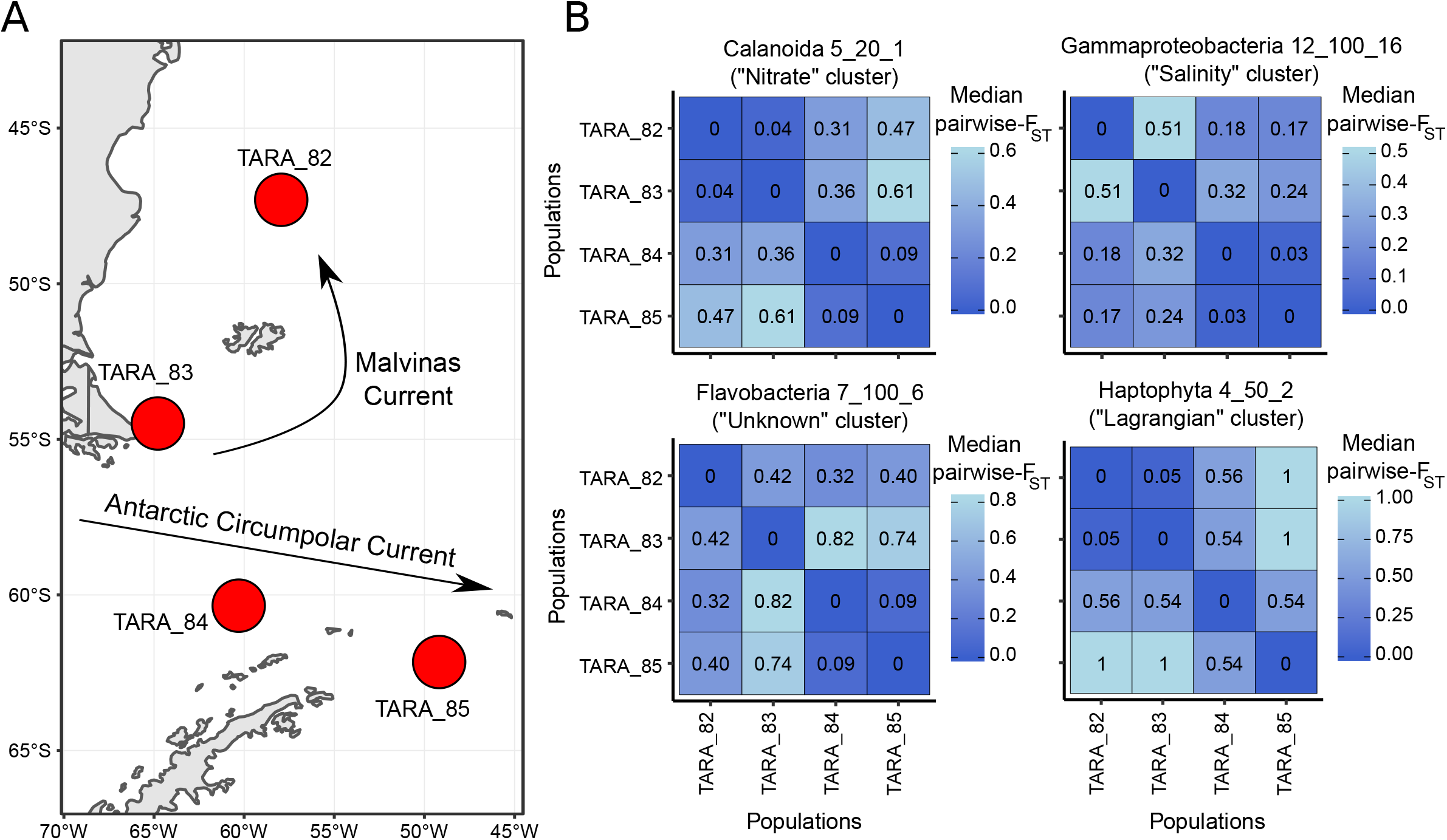
Genomic differentiation in Southern Ocean. A) Map localizing TARA_82, 83, 84, 85. The two arrows correspond to the trajectories of currents, based on Lagrangian trajectories, travel times and literature B) Pairwise-*F_ST_* matrices of the four MVSs specific to this area.

## Discussion

### Metavariant species as a representation of species polymorphism

Metavariant species were detected in each of the four size fractions. The number of genomic variants varied from 114 to several hundreds, with a very low rate of estimated false positive metavariants (46) enabling a realistic overview of the population structures of marine planktonic species lacking reference sequences. With this approach, metagenomic data help us to draw the silhouette of species population structure while previous studies are often based on few genetic markers, few samples, and are restrained to small geographic areas.

We were able to detect an extensive range of taxa, reflecting the biodiversity of epipelagic layer of oceans. It must be noticed that for each MVS, a majority of variant sequences didn’t show any taxonomic signal, an observation already made in other studies using *Tara* Oceans data (50,51). The level and quality of taxonomical assignation are both due to a lack of references in databases and to the small size of the sequences, reducing the chance of matching a reference and having a clear assignation.

Notwithstanding these technical limits for the taxonomical annotation of the MVS, four notable taxonomic groups retrieved from MVSs can be described and be related to previous observations. First, we were able to detect a virus, from the order of Caudovirales, and probably belonging to the bacteriophage family of Myoviridae. These viruses are known to be abundant compared to other viruses in oceans (75)◻, notably infecting Cyanobacteria (i.e. Prochlorococcus and Synechococcus), and constitute the majority of viral populations in GOV 2.0 (76)◻. Second, two Cyanobacteria, probably two Synechoccocus (15_500_9 and 7_20_37) were detected in the same locations in Mediterranean Sea, with clear *F_ST_* unimodal distributions (Supplementary Figure S6) and could be related to already observed ecotypes of Mediterranean Synechoccocus (77)◻. Third, in protists, two MVSs corresponding to Mamiellales (6_5_14 and 9_500_10) are respectively located in *Tara* stations where *Bathycoccus prasinos* and *Bathycoccus spp. TOSAG39–1* were the most abundant (Supplementary Table S2) in a previous study using *Tara* Oceans metagenomic dataset (78)◻. Finally, copepods formed the largest group retrieved by metaVaR, with a predominance of calanoid species compared to cyclopoid species. Finding a high number of these species was expected, considering copepods are very abundant in oceans (79,80) and well represented in the *Tara* Oceans dataset (51).

Together, these MVSs show the ability of metaVaR and our taxonomic assignation to distinguish closely-related species or ecotypes, and the accuracy to retrieve species.

### Differentiation of plankton populations from a global view

Our results showed clear patterns of differentiation among MVSs that depend on the basins and the size of organisms. Populations belonging to different basins tend to be more differentiated than populations located in the same basins, which could be explained by relatively smaller connections within basins than between basins. While this trend has been observed several times (28,81,82)◻, it hides interesting patterns. We observed the central place of NAO, relatively well connected to both MED and SAO, and a slightly lower connection between MED and SAO. Also, the SO was characterized by a relative isolation from the other basins. Indeed, SO shares few MVSs with other basins, and the latter are relatively highly differentiated. This situation was already observed notably in the copepod *Metridia lucens* (83)◻, with important differences between the populations of the basin. This area is characterized by differences in environmental conditions among it, and compared to the rest of the basins, with higher silicate, nitrate and phosphate concentrations on one hand, and lower salinity and temperature on the other hand (Figure 4B, Supplementary Figure S3). Plus, water masses are driven over thousands of kilometres by the complex Antarctic Circumpolar Current (ACC) (84)◻, which could favour gene flow between long-range locations all around the Antarctic. In addition, the Lagrangian data clearly traced the northward Malvinas current (an ACC branch), which mixes hot waters from the Brazil current with cold waters of the ACC in the Brazil–Malvinas Confluence(85)◻, possibly favouring the isolation of species in the south of this area. This situation could explain why these MVSs are both specific to Austral *Tara* stations and highly differentiated.

We showed that smaller organisms, like protists and bacteria, are more structured throughout oceans than zooplankton. These groups are not characterized by the same range of population sizes, dispersal capacities nor generation times, leading to different effects on their evolution. Finally, we were able to see a unique diversity of population differentiation among MVSs (Figure 3E), from unstructured to highly differentiated MVSs. The latter observation could be understood as the capacity of MVSs to capture complexes of closely-related species, as already described, for example, in *Oithona similis* in the NAO, SAO, SO and Arctic Ocean (86).

However, limitations arise from the use of *F_ST_*, which is affected by population effective size, described as high in plankton organisms in the few studies that estimated this parameter (13,87,88).

### Lagrangian travel times to estimate marine current transport

From the computation of Lagrangian travel times and sampling sites clustering, we were first able to distinguish three basins: NAO, MED and SAO-SO. Interestingly, the isolation of SO is not observed here, reinforcing our previous observations of genetic specificities linked to the unique environmental conditions of this basin. However, important differences were also observed between and among basins. For example, the Eastern part of the SAO presented an important connection with the NAO, which reflects the North Equatorial Current that linked these locations. Moreover, we saw how travel times from the SO to the Eastern part of the SAO were relatively small, which we can be linked to the Antarctic Circumpolar Current. Inside NAO, travel times between *Tara* oceans sampling sites presented a clear West-East trend, with some local divergences, which is related to the Gulf Stream and the North Atlantic Drift. Finally, inside MED, we clearly observed a West-East trend, with three different patterns: TARA_7/9/11 in the Western basin, TARA_18 to 26 in the Eastern basin, and the relative isolation of TARA_30 in the Levantine part of MED. Finally, the Haptophyta MVS from SO presented a differentiation linked to both the ACC and the Malvinas Current with the populations of TARA_83 and TARA_82 being highly connected by the fast Malvinas Current and a progressive eastward increase of *F_ST_* from TARA_83, TARA_84 and TARA_85.

Altogether, these results show the accuracy of this computation to reflect some of the main surface marine currents and the connectivity between *Tara* stations.

### Shaping of genomic differentiation by marine currents and environmental factors

In this study, the genomic differentiation of planktonic species was partially linked to environmental parameters and Lagrangian travel times. We first saw that globally, marine currents, salinity and temperature were the most important tested drivers of genomic differentiation, and that nitrate, silicate and phosphate had a relatively lower impact and this does not seem to be clade specific. Salinity and temperature are known to affect biogeography, community composition and population structure (15,28,43,49,89)◻. The role of nutrients like nitrate (90)◻, silicate (25,91,92)◻ and phosphate (93)◻ in marine micro-organisms metabolism, diversity and in the frame of their biogeochemical cycles (94–96) has been well studied, but their impact on the population structure has never been investigated at this scale.

This study also points to the importance of computing Lagrangian travel time estimates to evaluate the role of transport by marine currents, that is critical for the understanding of plankton genomic differentiation, as underlined here and in previous studies (34,36,40,97). We can note that obtaining proper haplotypes or genotypes together with considering the asymmetric travel times between locations would allow measuring the directional gene flow between populations.

We also notice that a large part of genomic differentiation cannot be explained in this study. The absence of physico-chemical parameters like metals, a key for cellular metabolism (19,98), sulfur (99)◻ or pH (18) could also enhance our comprehension of plankton genomic differentiation. Also, the contribution of biotic interactions between trophic levels, like grazing on phytoplankton by zooplankton (100)◻ should also be examined.

### Plankton connectivity as a mosaic

Finally, in our study, the identification of group of planktonic species having similar genomic differentiation trends driven by abiotic factors clearly demonstrated the mosaic of plankton population differentiation. This mosaic trend is underlined by the diversity of environmental conditions influencing the differentiation but was also exampled by the absence of link between the number of basins where MVSs were detected and their global differentiation (Figure 3F) and with several individual cases. This shows that the living range of species is not correlated to their population structure, i.e. cosmopolitan species do not necessarily present an absence of population structure and species with populations present in close locations can exhibit high differentiation (such as SO). We thus showed how population genomics is important to decipher the connectivity of plankton, and can be complementary to the traditional metabarcoding approach, that fails to quantify the connectivity and intra-species structure patterns.

Furthermore, we showed that the clade of species was not determinant to identify the drivers of the genomic differentiation.

The next step would be to better catch the relative effects of evolutive forces on genome, like genetic drift and selection, as the question is still unresolved (13,15–17)◻. Sequencing genomes or haplotypes data could resolve this question, but in the frame of metagenomic, the latter is still a technical and computational challenge.

## Acknowledgments

We thank the Commissariat à l’Energie Atomique et aux énergies altenatives, France Génomique (ANR-10-INBS-09), and Oceanomics (ANR-11-BTBR-0008). We acknowledge Paul Frémont for his help with WOA environmental parameters. This is contribution number XX from *Tara* Oceans.

## Author’s contributions

RLJ performed all analyses. MAM designed and supervised the study. CA gave expertise support on the statistical framework. MO computed Lagrangian travel time estimates and MO and AS offered expertise on these results. PW offered scientific support.

## Data availability

The set of MVSs is available on github at: https://github.com/rlasojad/Metavariant-Species

## Competing interests

The authors declare no competing interests.

## Supplementary Tables

**Supplementary Table S1: Summary of MVSs**

**Supplementary Table S2: MVSs and *Bathycoccus***

## Supplementary Figures

**Supplementary Figure S1 : MetavaR clustering**

**Supplementary Figure S2 : Overview of taxonomic assignation**

**Supplementary Figure S3 : Environmental parameters maps**

**Supplementary Figure S4 : Principal component analysis of the contribution of environmental parameters to the genomic differentiation of MVSs**

**Supplementary Figure S5 : Occurrence of MVSs**

**Supplementary Figure S6 : Global distributions of *F_ST_***

**Supplementary Figure S7 : Lagrangian estimates matrices**

**Supplementary Figure S8: Lagrangian trajectories for stations of Southern Ocean.**

## Notes

### Competing Interest Statement

The authors have declared no competing interest.

