## Supplementary Figures for "How marine currents and environment shape plankton genomic differentiation: a mosaic view from *Tara* Oceans metagenomic data"

Table of contents

Supplementary Figure S1: MetaVaR clustering	3

Supplementary Figure S2: Overview of taxonomic assignation	4

Supplementary Figure S3: Environmental parameters maps	5

Supplementary Figure S4: Principal component analysis of the contribution of environmental parameters to the genomic differentiation of MVSs	8

Supplementary Figure S5: Occurrence of MVSs	9

Supplementary Figure S6: Global distributions of *F_ST_*	10

Supplementary Figure S7: Lagrangian estimates matrices	11

Supplementary Figure S8: Lagrangian trajectories for stations of Southern Ocean.	12

Supplementary Table S2: MVSs and *Bathycoccus*	15

### Supplementary Figure S1: MetaVaR clustering

Number of MVCs found for each dataset, and for each couple of dbscsan parameters ε and minimum points (MinPts). In blank, no cluster were found for the corresponding parameters.

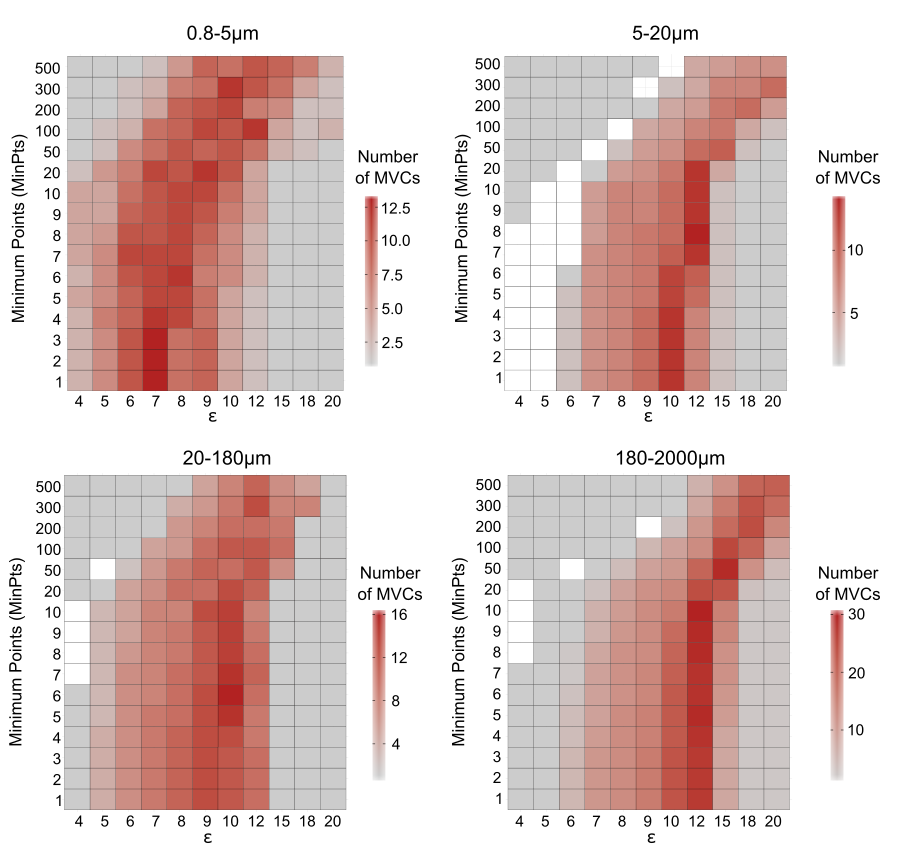

### Supplementary Figure S2: Overview of taxonomic assignation

Pipeline describing how each MVS was assigned to a taxonomic group.

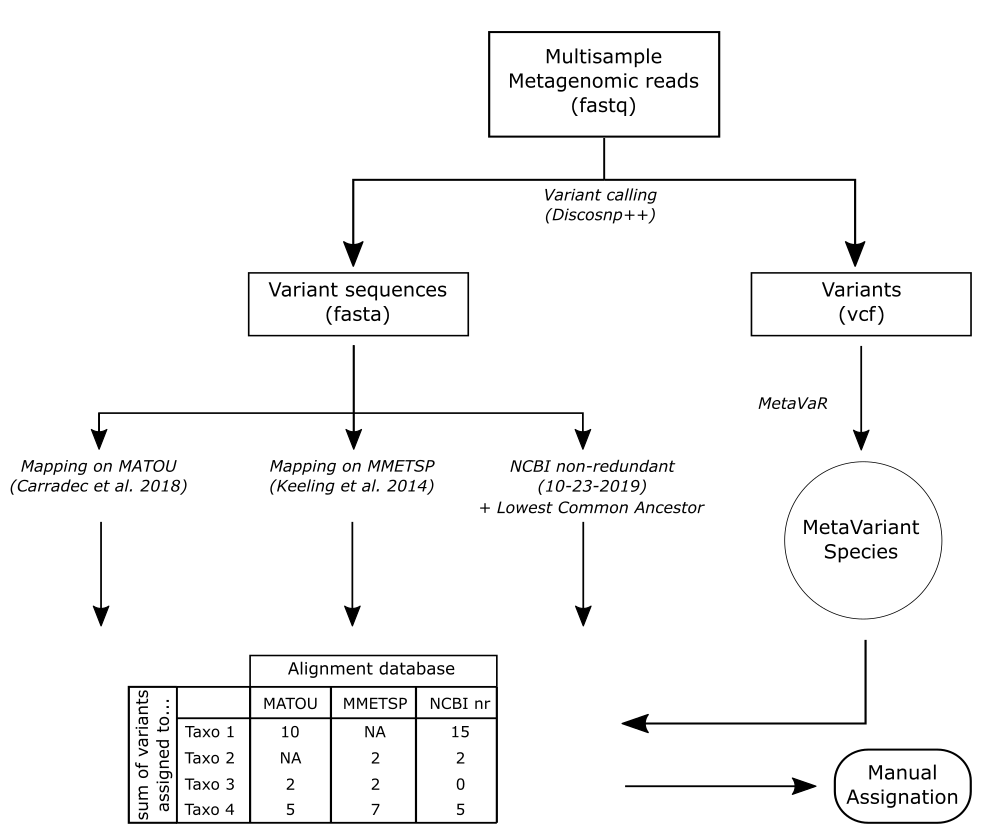

### Supplementary Figure S3: Environmental parameters maps

Each dot corresponds to a *Tara* station.

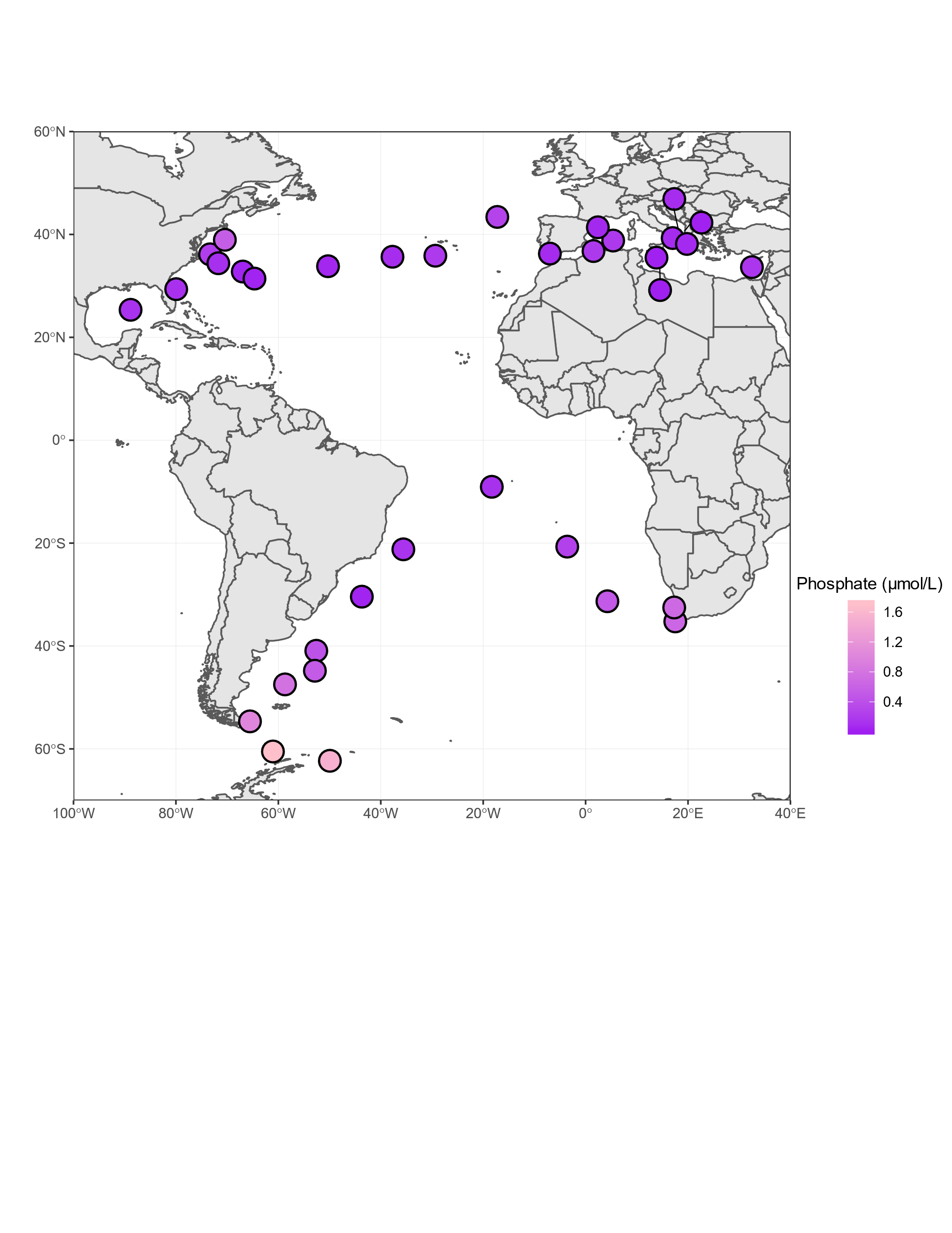

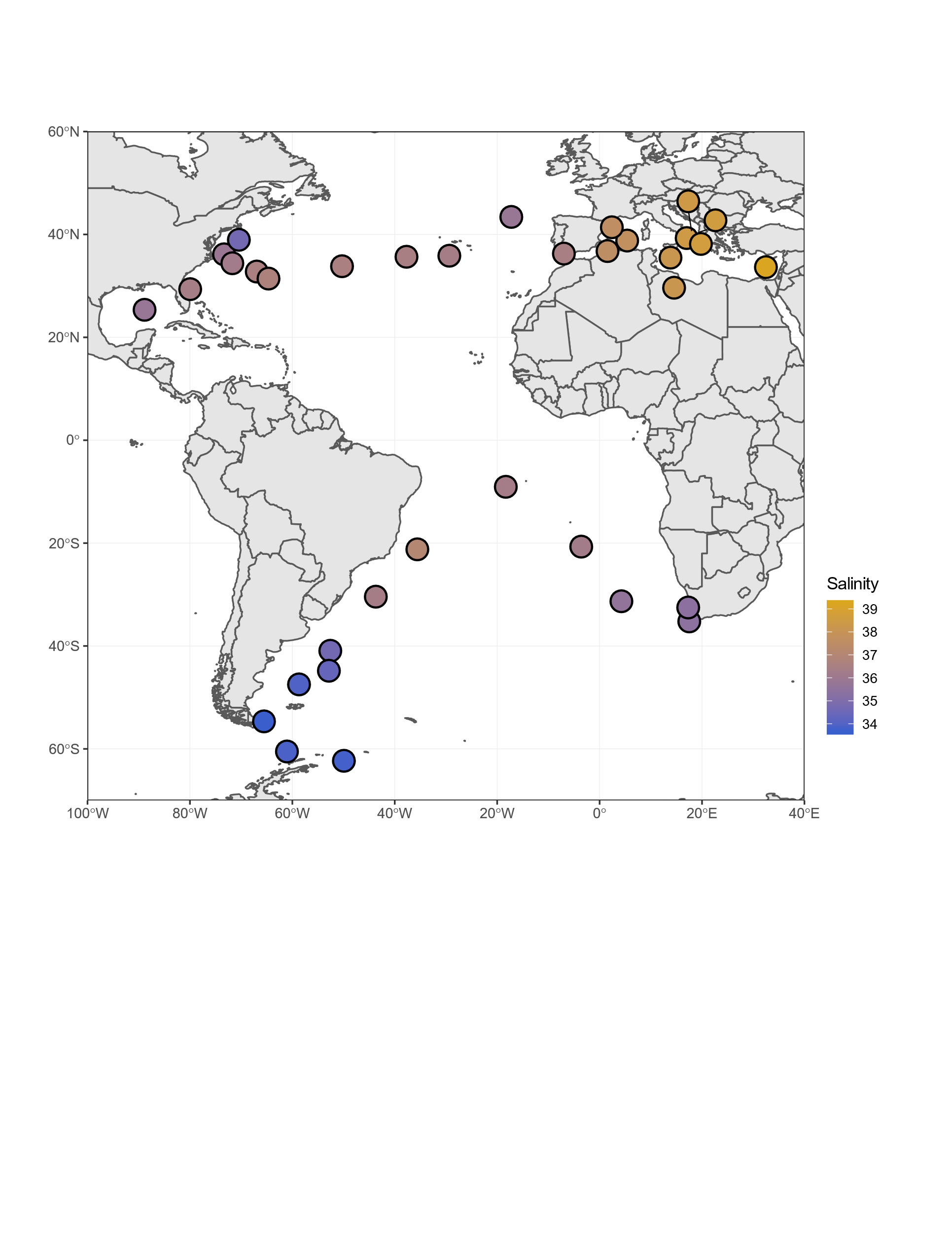

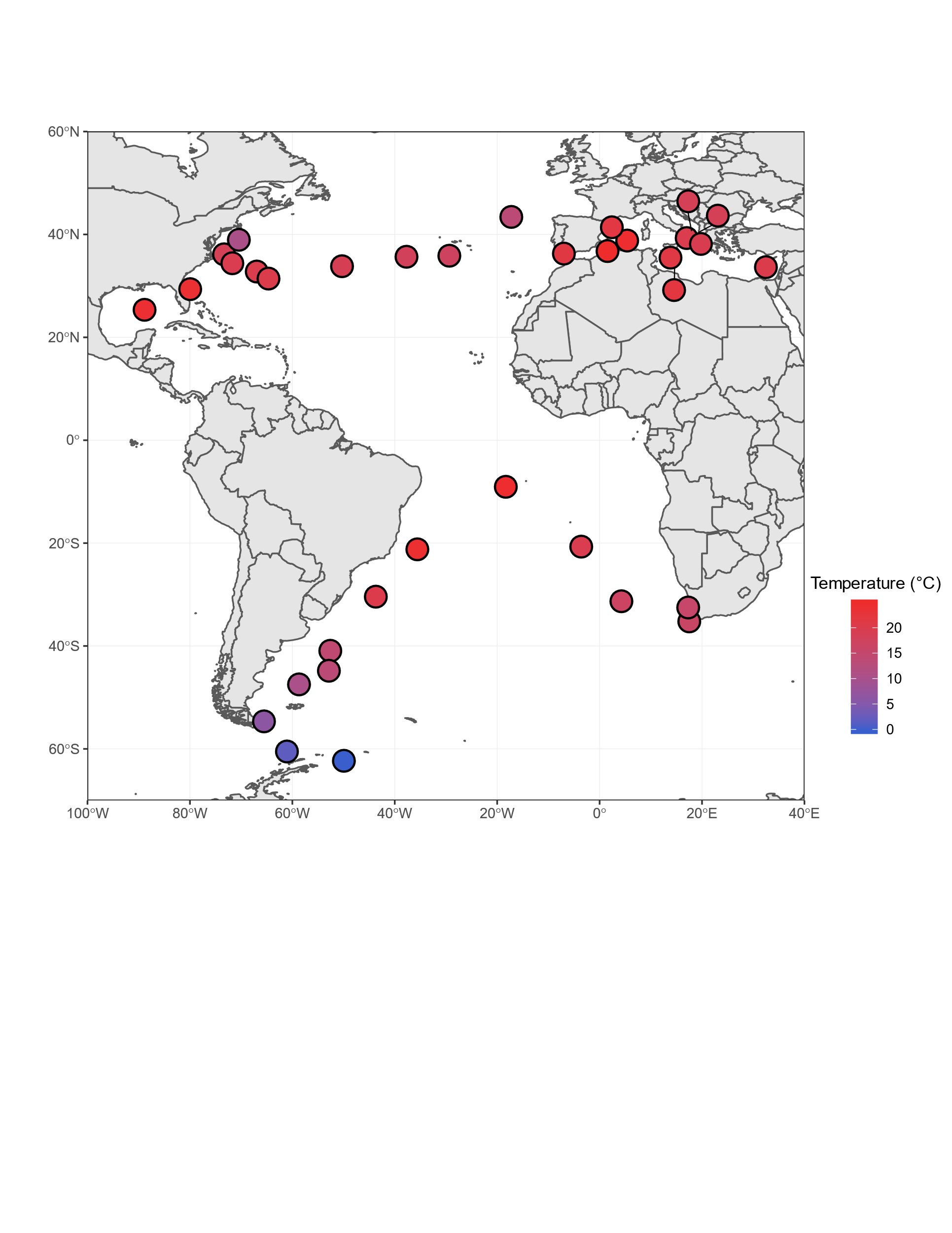

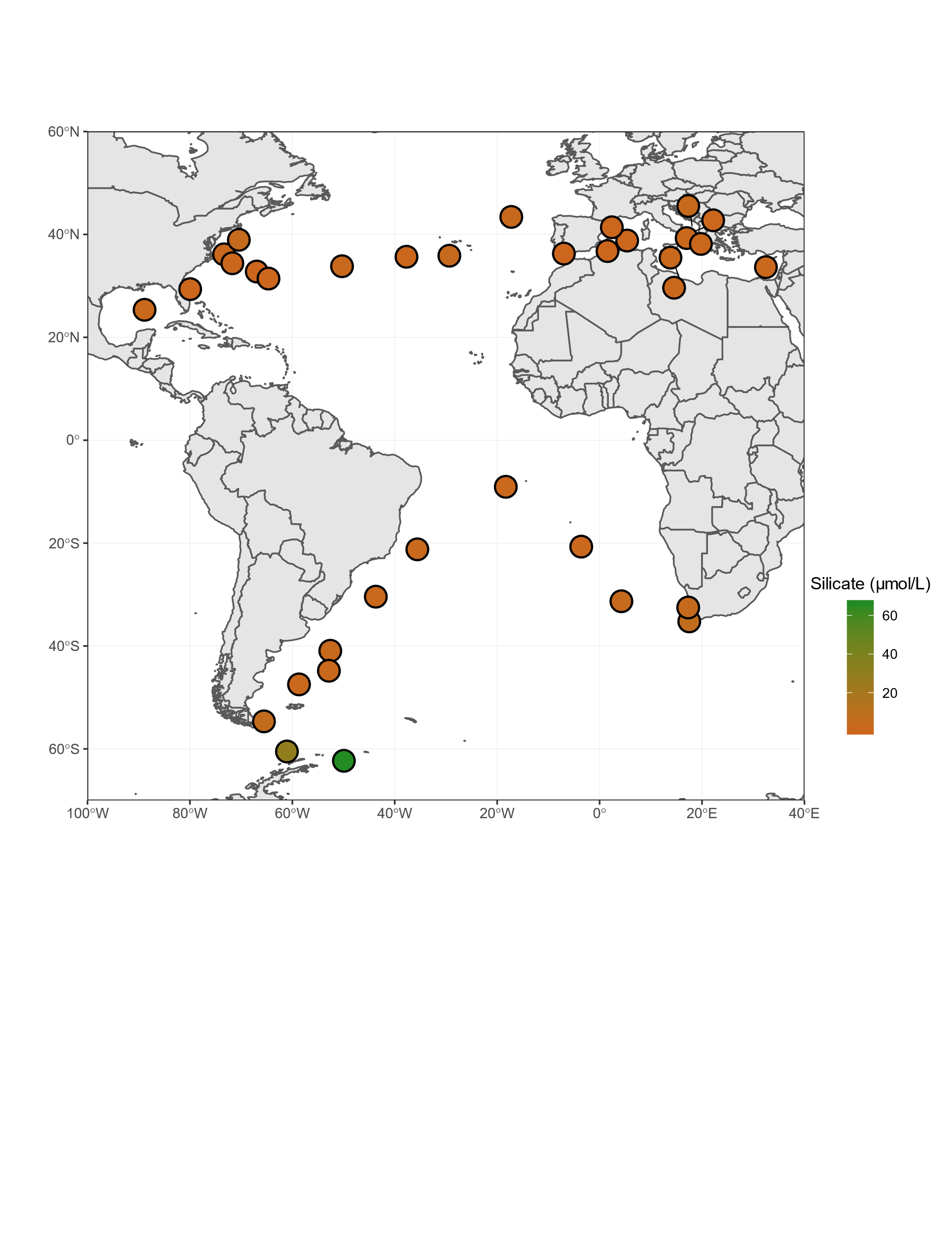

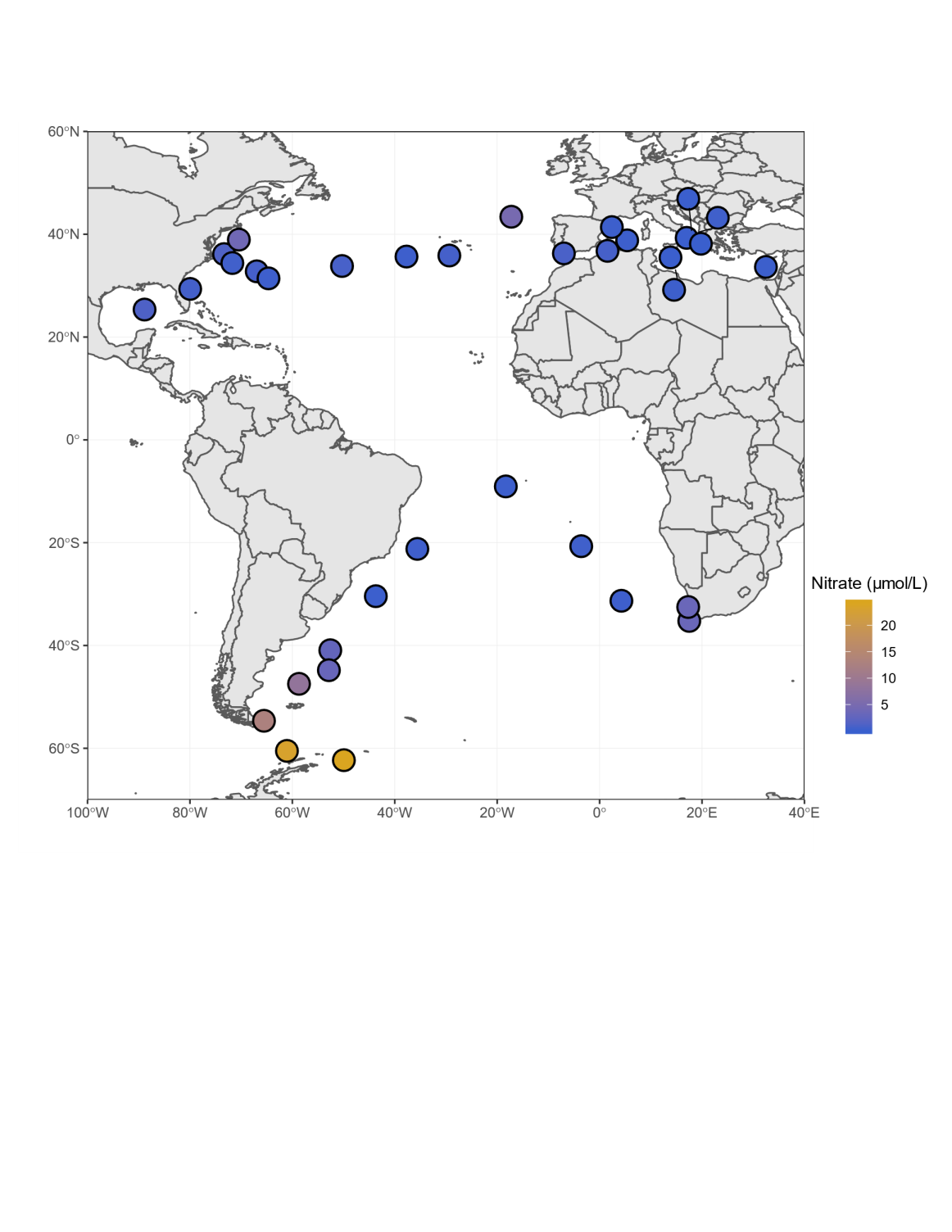

### Supplementary Figure S4: Principal component analysis of the contribution of environmental parameters to the genomic differentiation of MVSs

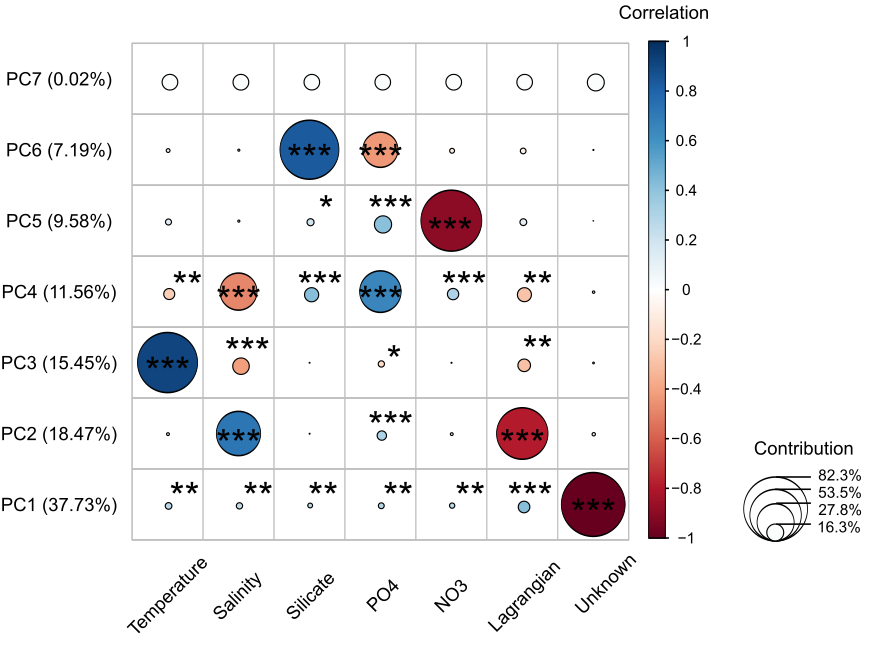

### Supplementary Figure S5: Occurrence of MVSs

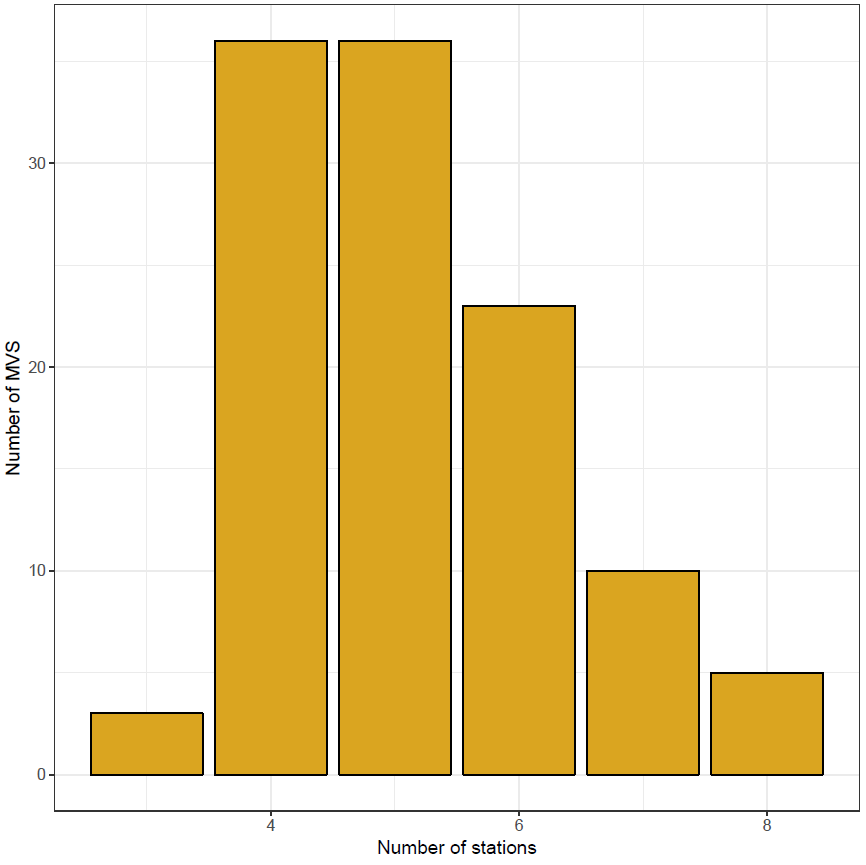

### Supplementary Figure S6: Global distributions of *F_ST_*

Each plot corresponds to an MVS. The color of the violin is linked to the taxonomy, and the background color of MVSs’ names stand for the size fractions; red, blue, green and yellow for 0.8-5µm, 5-20µm, 20-180µm and 180-2000µm respectively.

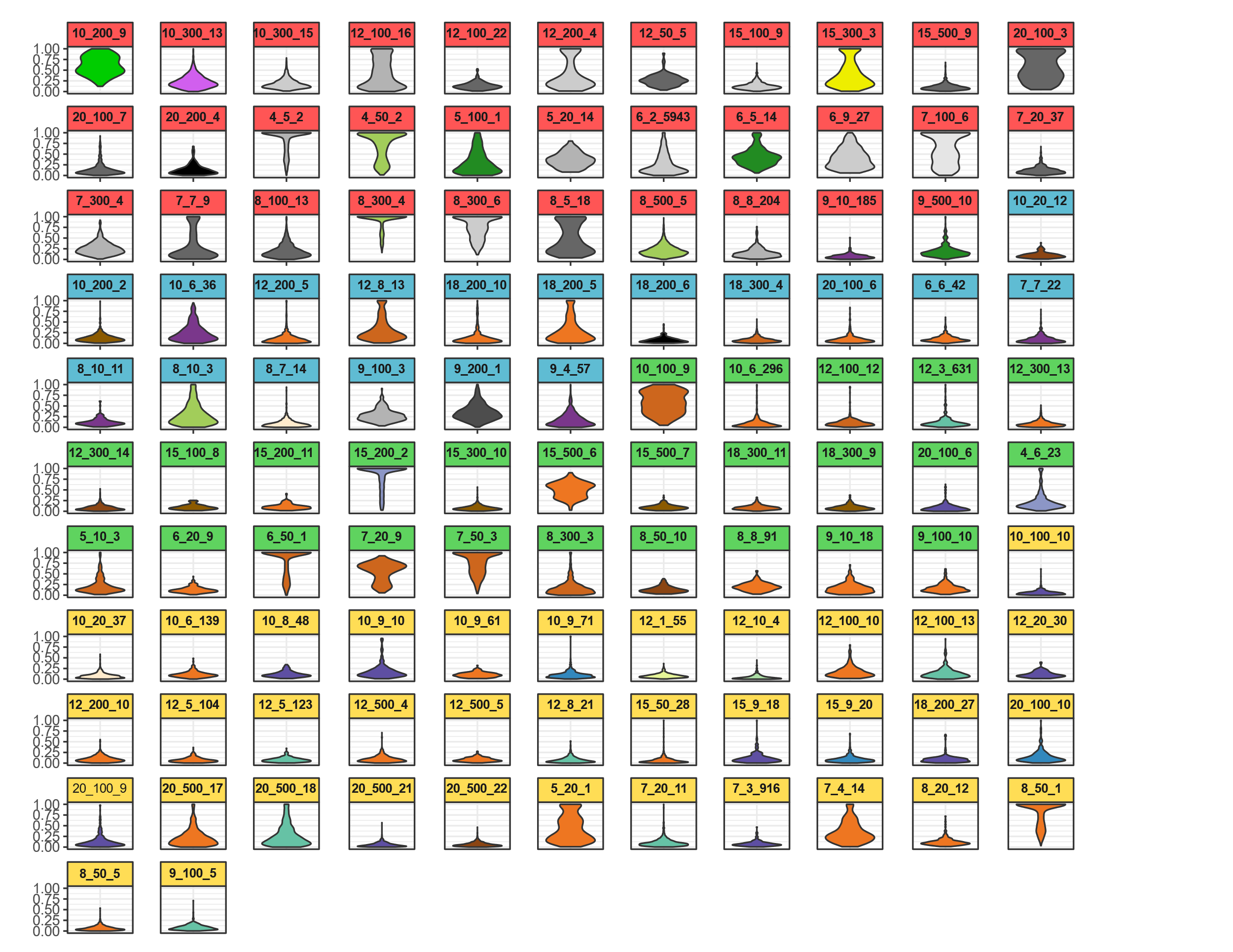

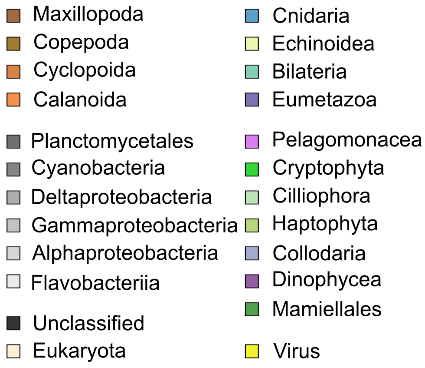

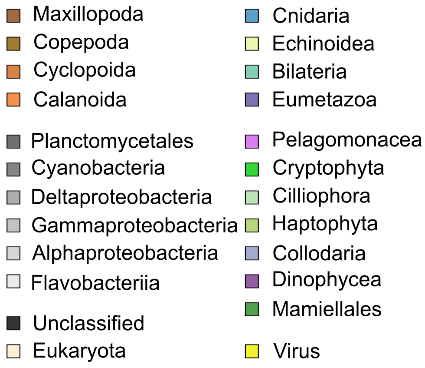

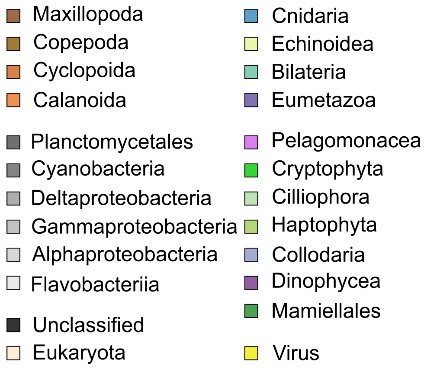

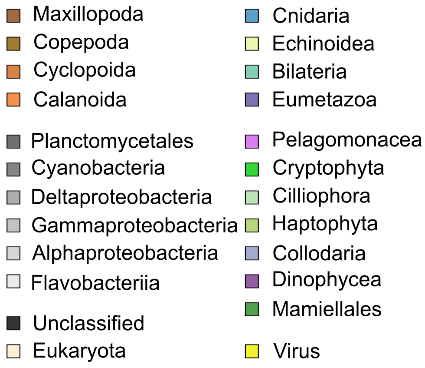

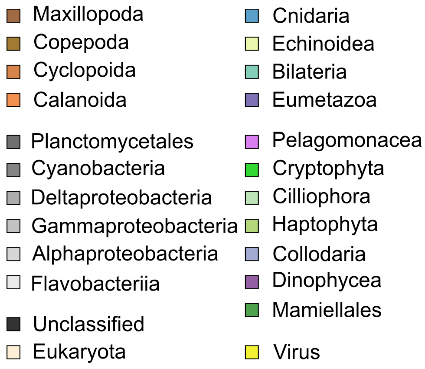

### Supplementary Figure S7: Lagrangian estimates matrices

Results of Lagrangian travel time computations. A) Asymmetric times between the 35 stations. Because of the important difference in travel times between Mediterranean Sea stations and the rest, we also present the Lagrangian estimates between B) Mediterranean Sea stations, C) Atlantic and Southern Oceans stations.

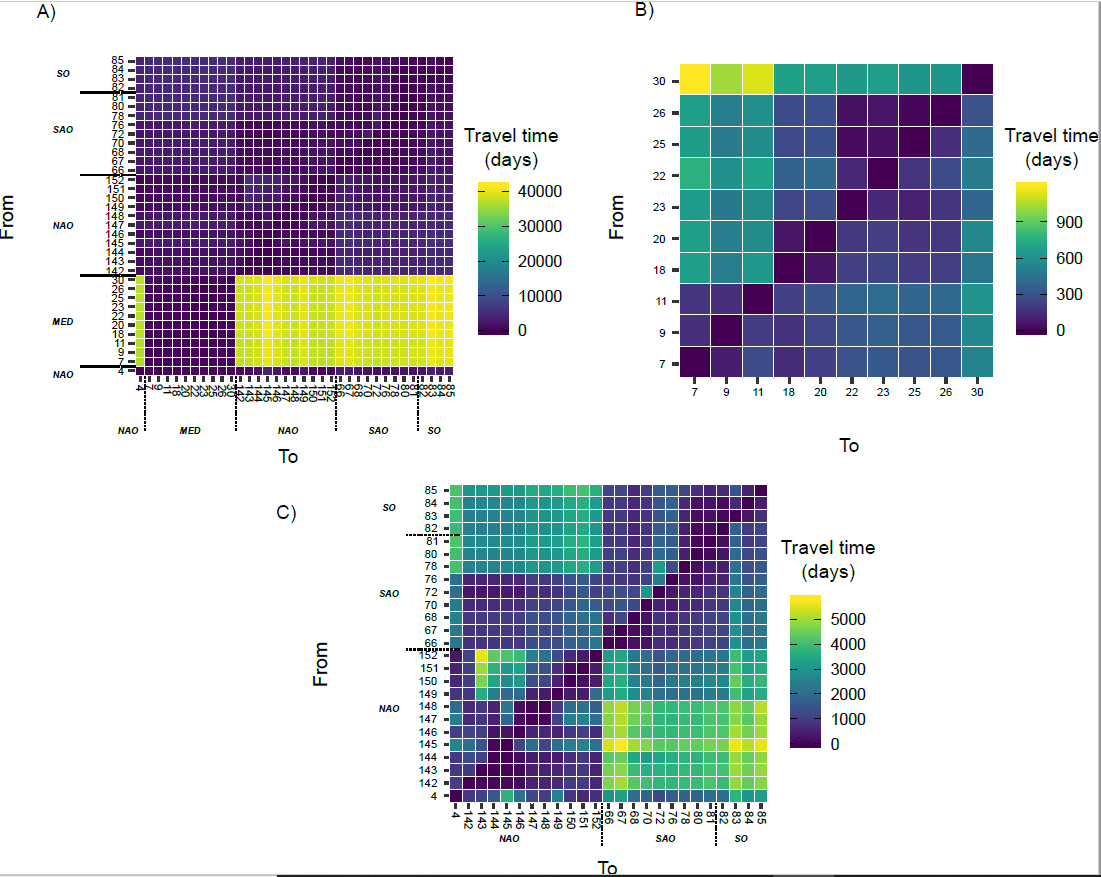

### Supplementary Figure S8: Lagrangian trajectories for stations of Southern Ocean.

For each pair of stations; the two upper plots are the drifters trajectories with the fastest and the slowest tracks in blue and red respectively and the lower plots picture the distribution of Lagrangian travel times estimated after bootstrap for the corresponding trajectory.

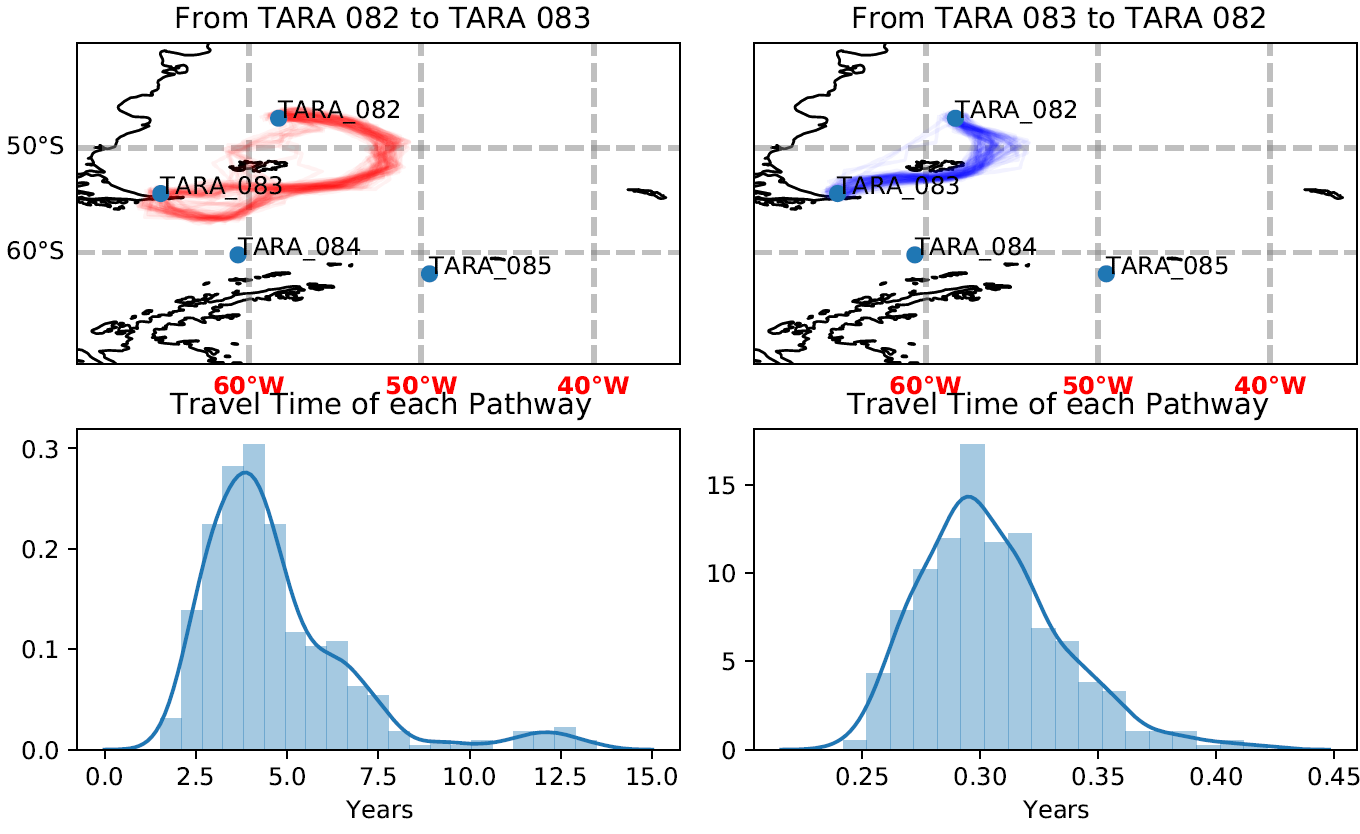

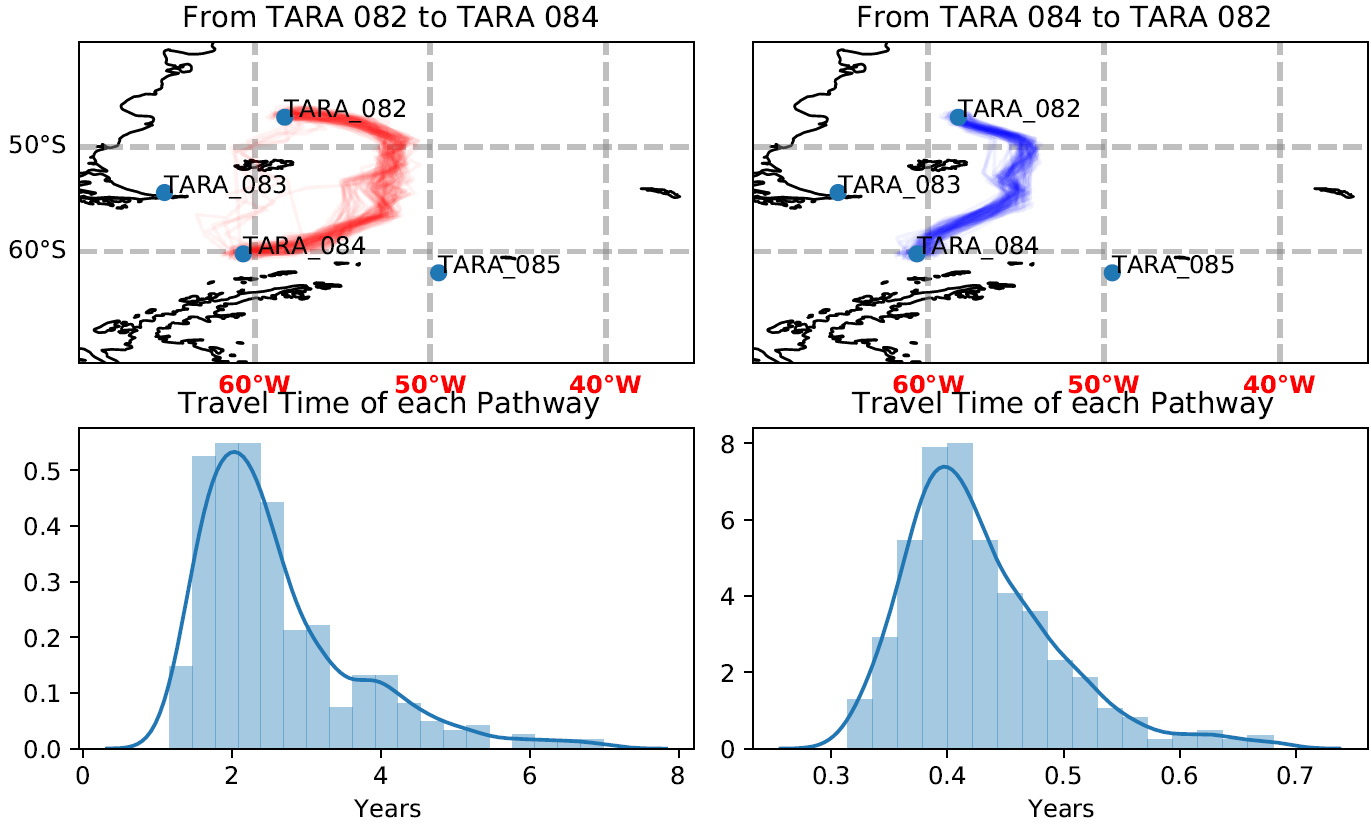

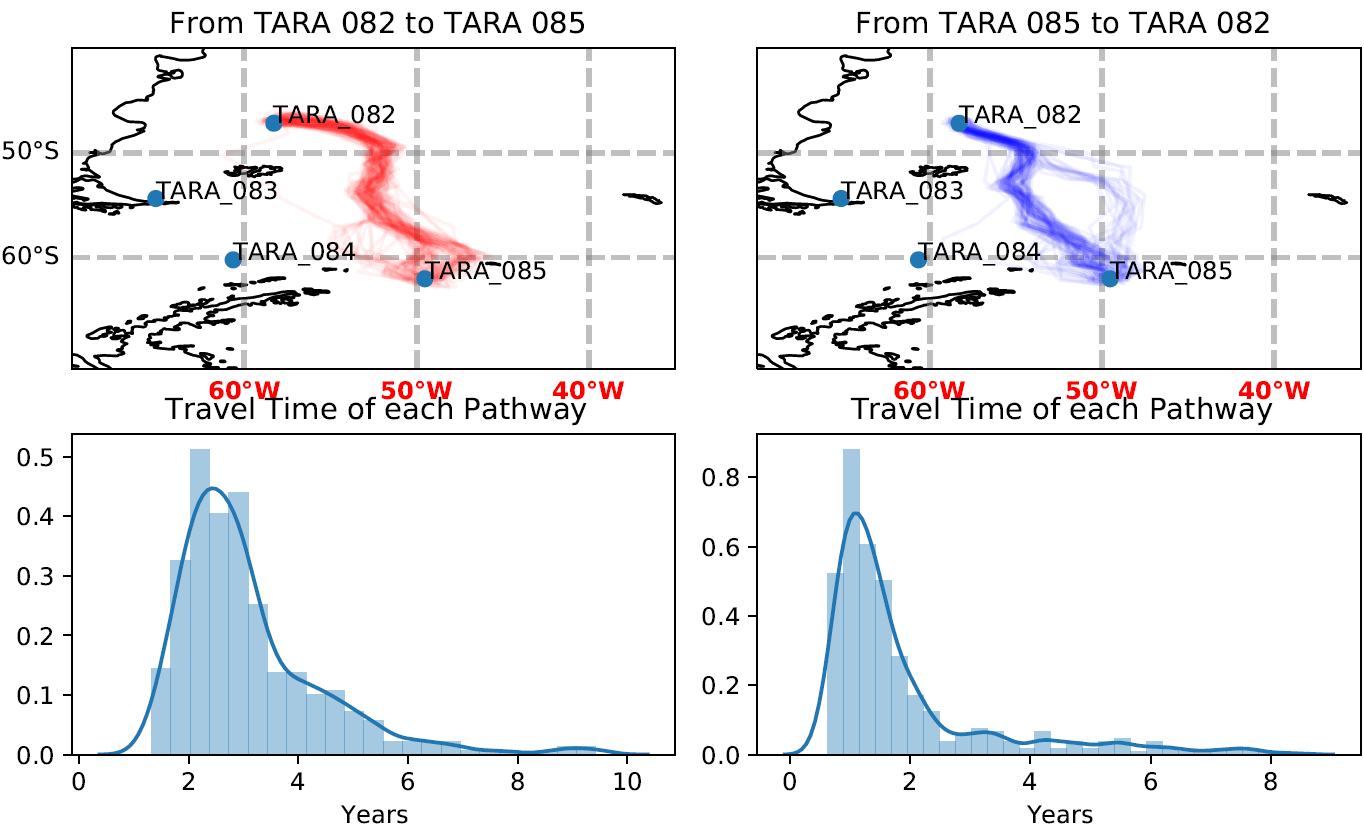

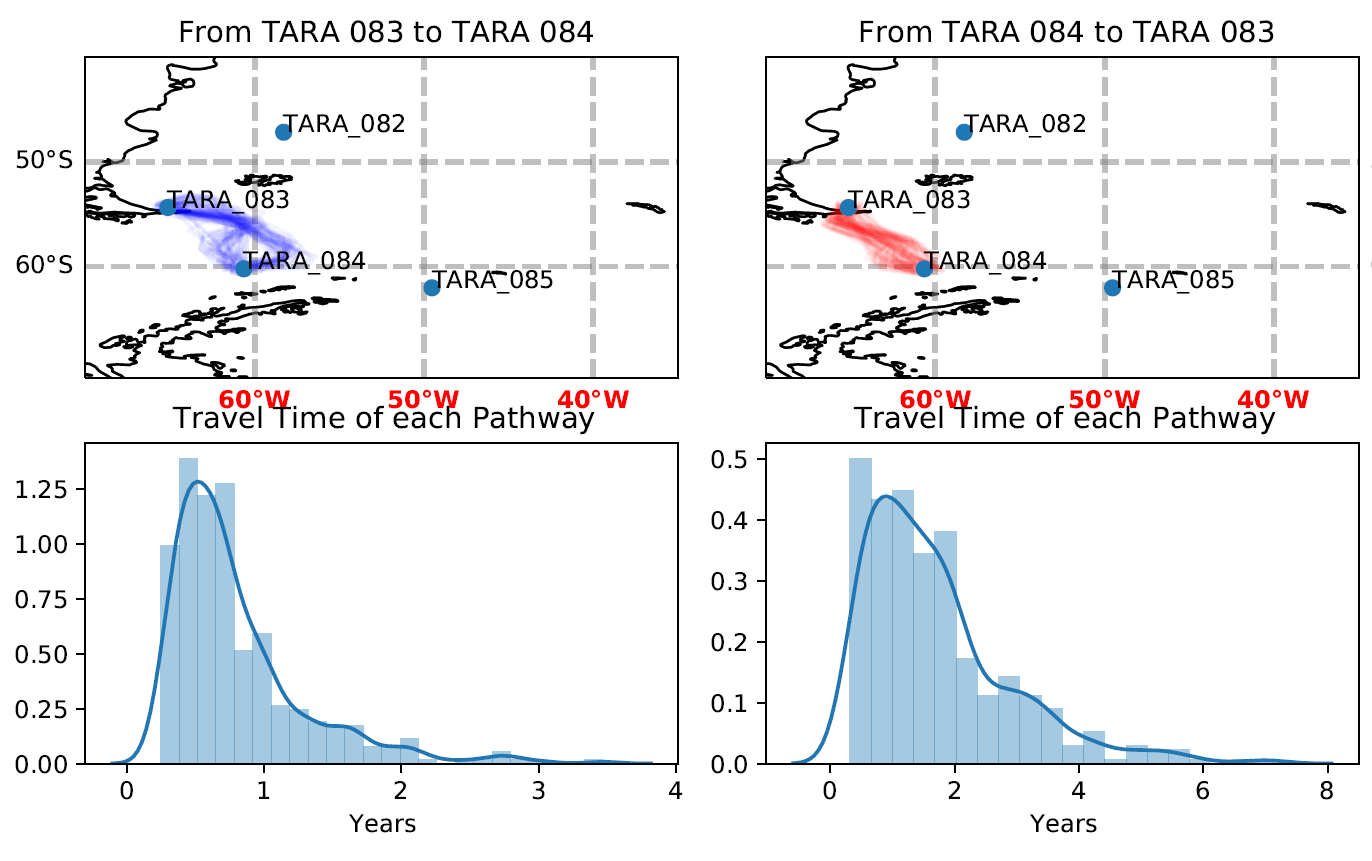

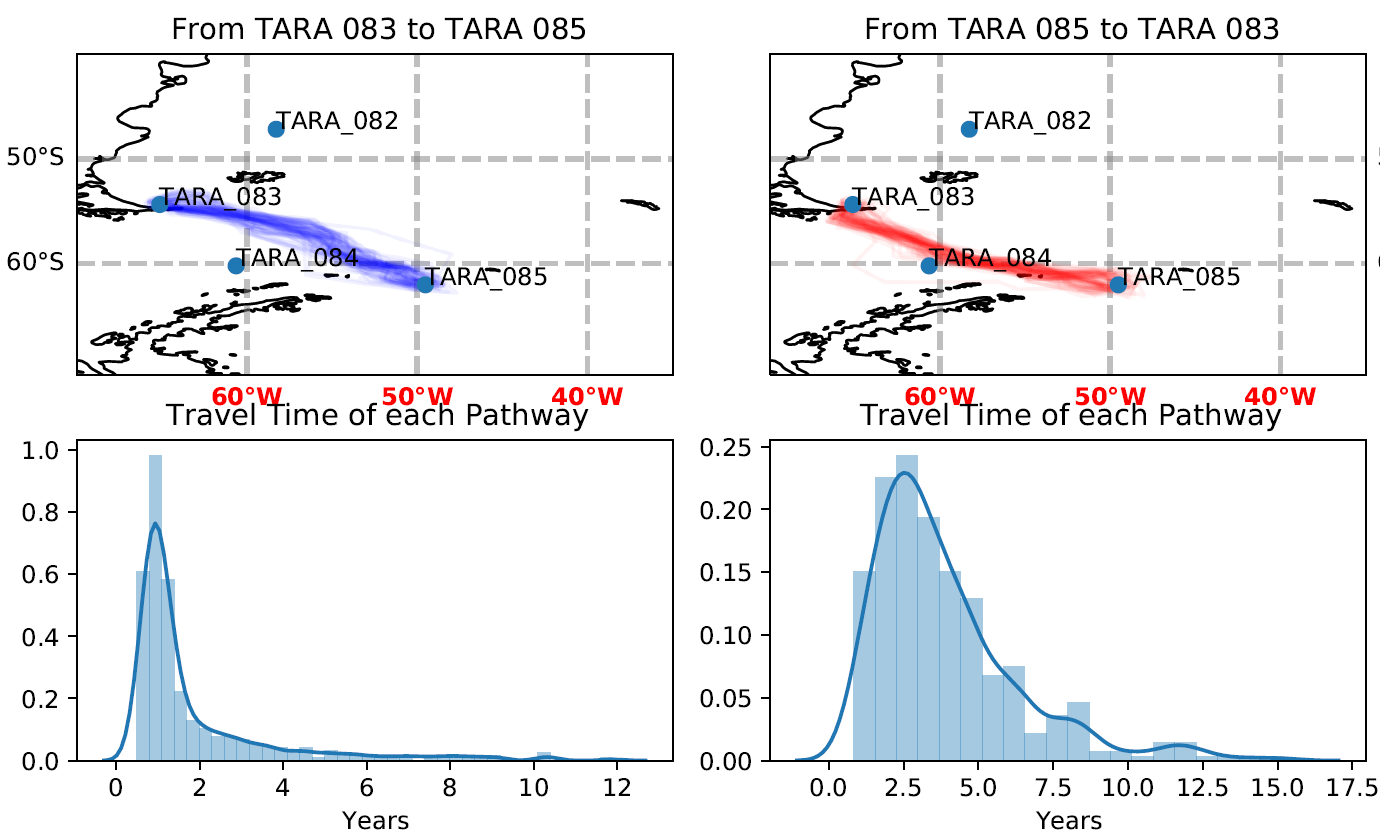

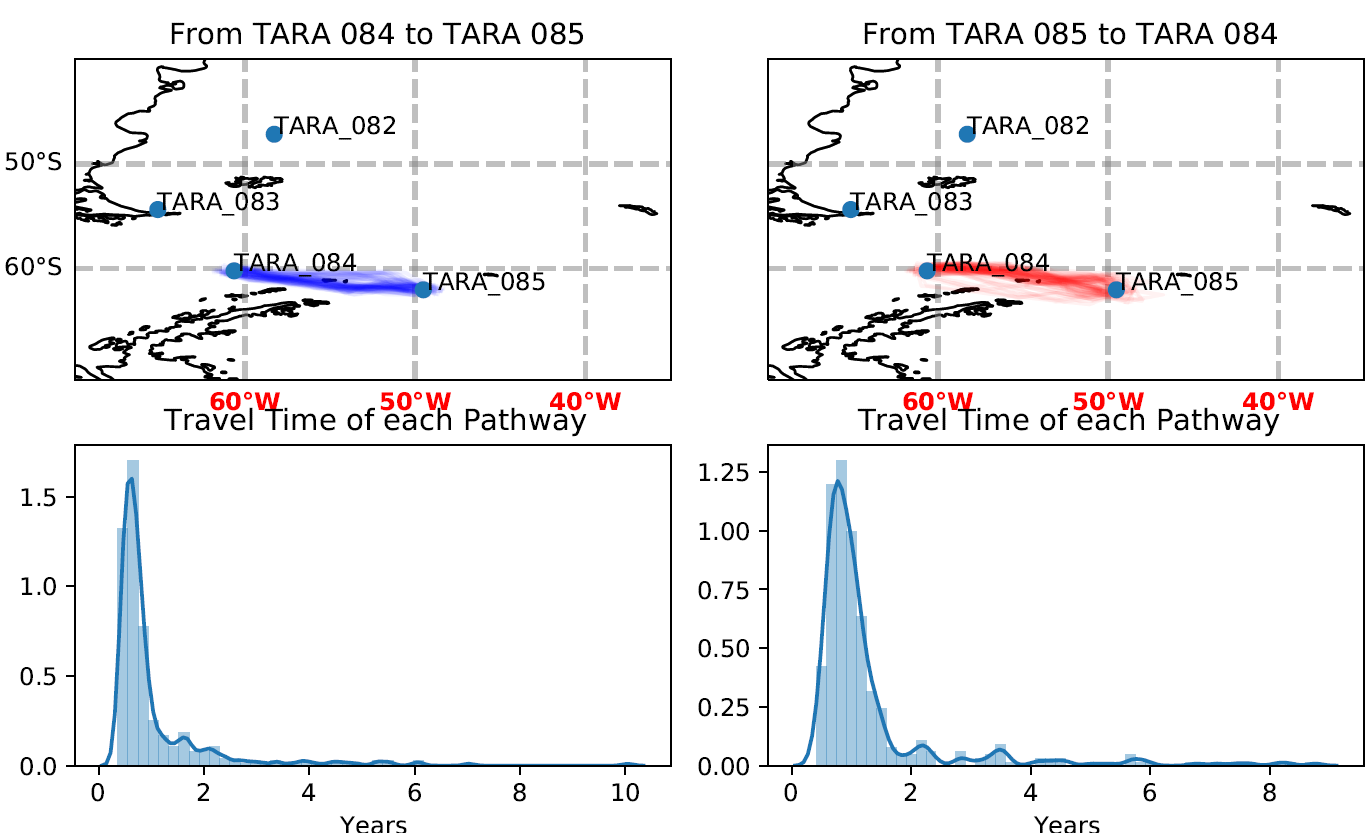

### Supplementary Table S2: MVSs and *Bathycoccus*

The columns *"Bathycoccus"* MVSs reflects the occurrences of the two MVSs identified as potential *Bathycoccus* in our dataset. The columns “*Bathycoccus* strains” are the percentage of metagenomic reads from each *Tara* stations matching the two reference genomes (data extracted from Leconte et al. 2020). MVSs 6_5_14 and 9_500_10 are present where *Bathycoccusprasinos* RCC1105 and *Bathycoccus* TOSAG39.1 are the most abundant, respectively.

|  | *"Bathycoccus"* MVSs | | *Bathycoccus* strains | |
| --- | --- | --- | --- | --- |
| *Tara* stations | 6_5_14 | 9_500_10 | *Bathycoccus prasinos RCC1105* | *Bathycoccus TOSAG39.1* |
| TARA_66 | Yes | No | 0.7600 | 0.1264 |
| TARA_67 | Yes | No | 0.9098 | 0.0156 |
| TARA_80 | Yes | Yes | 0.9215 | 0.3204 |
| TARA_81 | Yes | No | 1.3416 | 0.0202 |
| TARA_142 | No | No | 0.0005 | 0.0345 |
| TARA_145 | Yes | No | 1.3493 | 0.1263 |
| TARA_146 | No | Yes | 0.1010 | 1.8254 |
| TARA_147 | No | Yes | 0.0906 | 0.8468 |
| TARA_150 | No | Yes | 0.3085 | 0.2685 |
| TARA_152 | Yes | No | 0.4797 | 0.0329 |
