## Supplementary Table S1 for "How marine currents and environment shape plankton genomic differentiation: a mosaic view from *Tara* Oceans metagenomic data"

| MVS name | Size Fraction | Taxonomy (precise) | Retained Taxonomy | Larger group | Number of variants (MVC) | Number of variants (MVS) | Basin(s) | Tara stations |
|---|---|---|---|---|---|---|---|---|
| 15_200_2 | 20-180µm | Ciliophora (Spirotrichea) | Ciliophora | Unicellular Eukaryotes | 1363 | 695 | MED-NAO-SAO | TARA_145,23,66,70 |
| 15_300_10 | 20-180µm | Maxillopoda (Calanoid) | Maxillopoda | Copepods | 1237 | 715 | MED-NAO-SAO | TARA_150,22,25,26,70 |
| 15_500_6 | 20-180µm | Copepod (Calanoid) | Calanoida | Copepods | 1006 | 221 | MED-NAO | TARA_144,149,150,18,23 |
| 15_500_7 | 20-180µm | Maxillopoda (Cyclopoid) | Maxillopoda | Copepods | 2274 | 346 | NAO-SAO | TARA_144,146,147,148,149,70,76,78 |
| 18_300_11 | 20-180µm | Copepod (Cyclopoida) | Cyclopoida | Copepods | 1357 | 228 | NAO-SAO | TARA_143,144,146,147,149,150,70 |
| 18_300_9 | 20-180µm | Maxillopoda (Cyclopoid) | Maxillopoda | Copepods | 1770 | 665 | MED-NAO-SAO | TARA_72,143,26,70,76 |
| 20_100_6 | 20-180µm | Eumetazoa | Eumetazoa | Animals | 1019 | 474 | MED | TARA_7,11,22,23 |
| 4_6_23 | 20-180µm | Spirotrichea | Ciliophora | Unicellular Eukaryotes | 1092 | 711 | MED-SAO | TARA_23,25,26,78,80 |
| 5_10_3 | 20-180µm | Copepod (Oithona_nana?) | Cyclopoida | Copepods | 1434 | 113 | MED | TARA_7,11,22,23,26 |
| 6_20_9 | 20-180µm | Copepod (Calanoid) | Calanoida | Copepods | 1331 | 239 | NAO-SAO | TARA_72,142,143,144,148,76,78 |
| 6_50_1 | 20-180µm | Copepod (Cyclopoida) | Cyclopoida | Copepods | 1394 | 1266 | NAO-SAO | TARA_145,152,66 |
| 7_20_9 | 20-180µm | Copepod (Oithona) | Cyclopoida | Copepods | 1013 | 398 | MED-SAO | TARA_7,11,18,23,26,70 |
| 7_50_3 | 20-180µm | Copepod (Oithona) | Cyclopoida | Copepods | 1141 | 695 | NAO-SAO | TARA_148,150,67,68 |
| 8_300_3 | 20-180µm | Copepod (Oithona) | Cyclopoida | Copepods | 1229 | 912 | MED-SAO | TARA_7,18,26,70 |
| 8_50_10 | 20-180µm | Copepod (Cyclopoida) | Copepoda | Copepods | 2147 | 168 | MED-NAO | TARA_144,146,147,148,149,25,26,30 |
| 8_8_91 | 20-180µm | Copepod (Calanoid) | Calanoida | Copepods | 1234 | 150 | MED-NAO | TARA_144,147,150,151,25,30 |
| 9_10_18 | 20-180µm | Copepod (Calanoid) | Calanoida | Copepods | 1167 | 179 | MED-SAO | TARA_72,18,25,26,30,76 |
| 9_100_10 | 20-180µm | Copepod (Calanoid) | Calanoida | Copepods | 2127 | 227 | MED-NAO-SAO | TARA_72,143,151,7,18,30,70 |
| 10_100_10 | 180-2000µm | Eumetazoa | Eumetazoa | Animals | 1186 | 937 | NAO-SAO | TARA_72,142,143,144 |
| 10_20_37 | 180-2000µm | Eukaryota | Eukaryota | Poor classification | 1008 | 591 | NAO-SAO | TARA_147,148,149,150,70 |
| 10_6_139 | 180-2000µm | Copepod (Calanoid) | Calanoida | Copepods | 1017 | 238 | MED-NAO-SAO | TARA_149,150,151,152,9,25,68 |
| 10_8_48 | 180-2000µm | Eumetazoa | Eumetazoa | Animals | 1035 | 118 | NAO-SAO | TARA_147,148,149,150,151,4,78 |
| 10_9_10 | 180-2000µm | Eumetazoa | Eumetazoa | Animals | 1025 | 254 | MED-NAO-SAO | TARA_146,148,151,9,76,78 |
| 10_9_61 | 180-2000µm | Copepod (Calanoid) | Calanoida | Copepods | 1082 | 148 | NAO-SAO | TARA_142,143,144,150,151,70,76,80 |
| 10_9_71 | 180-2000µm | Cnidaria | Cnidaria | Animals | 1053 | 574 | MED-NAO | TARA_144,149,151,9,11,23 |
| 12_1_55 | 180-2000µm | Echinoidea | Echinoidea | Animals | 1094 | 535 | MED-NAO | TARA_142,144,146,147,148,149,30 |
| 12_10_4 | 180-2000µm | Collodaria | Collodaria | Unicellular Eukaryotes | 1068 | 962 | NAO | TARA_146,147,148 |
| 12_100_10 | 180-2000µm | Copepod (Calanoid) | Calanoida | Copepods | 1773 | 164 | MED-NAO-SAO | TARA_72,4,9,23,70,76 |
| 12_100_13 | 180-2000µm | Bilateria | Bilateria | Animals | 1237 | 466 | NAO-SAO | TARA_148,150,151,68,80 |
| 12_20_30 | 180-2000µm | Eumetazoa | Eumetazoa | Animals | 1067 | 122 | MED | TARA_9,11,20,22,23,25 |
| 12_200_10 | 180-2000µm | Copepod (Calanoid) | Calanoida | Copepods | 1099 | 753 | MED-NAO | TARA_148,149,150,151,25 |
| 12_5_104 | 180-2000µm | Copepod (Calanoid) | Calanoida | Copepods | 1006 | 422 | MED-NAO-SAO | TARA_144,150,20,25,68 |
| 12_5_123 | 180-2000µm | Bilateria (Chordata) | Bilateria | Animals | 1221 | 291 | MED-NAO | TARA_143,144,149,150,151,25 |
| 12_500_4 | 180-2000µm | Copepod (Calanoid) | Calanoida | Copepods | 1275 | 900 | MED-NAO | TARA_146,147,148,149,25 |
| 12_500_5 | 180-2000µm | Copepod (Calanoid) | Calanoida | Copepods | 2011 | 498 | MED-NAO-SAO | TARA_72,143,144,20,70,76 |
| 12_8_21 | 180-2000µm | Bilateria | Bilateria | Animals | 2749 | 1205 | NAO-SAO | TARA_72,142,143,76 |
| 15_50_28 | 180-2000µm | Copepod (Calanoid) | Calanoida | Copepods | 1020 | 845 | SAO | TARA_72,70,76,78 |
| 15_9_18 | 180-2000µm | Eumetazoa | Eumetazoa | Animals | 1007 | 390 | MED-NAO | TARA_143,144,20,22,25 |
| 15_9_20 | 180-2000µm | Cnidaria (Hexacorallia) | Cnidaria | Animals | 1121 | 426 | NAO | TARA_142,143,144,146,147,149 |
| 18_200_27 | 180-2000µm | Eukaryote (Eumetazoa) | Eumetazoa | Animals | 1113 | 233 | NAO-SAO | TARA_143,144,147,149,150,151,70 |
| 20_100_10 | 180-2000µm | Cnidaria (Hydrozoa) | Cnidaria | Animals | 1328 | 282 | NAO-SAO | TARA_72,143,4,76,80 |
| 20_100_9 | 180-2000µm | Eumetazoa | Eumetazoa | Animals | 1021 | 611 | MED-NAO | TARA_149,151,23,25,30 |
| 20_500_17 | 180-2000µm | Copepod (Calanoid) | Calanoida | Copepods | 1157 | 947 | MED-NAO | TARA_147,149,150,20 |
| 20_500_18 | 180-2000µm | Bilateria | Bilateria | Animals | 1280 | 1080 | MED-NAO-SAO | TARA_144,20,22,68 |
| 20_500_21 | 180-2000µm | Eumetazoa | Eumetazoa | Animals | 1102 | 832 | MED-NAO | TARA_147,149,150,20 |
| 20_500_22 | 180-2000µm | Copepod (Oithona) | Copepoda | Copepods | 1221 | 1007 | NAO | TARA_146,147,148,149 |
| 5_20_1 | 180-2000µm | Copepod (Calanoid) | Calanoida | Copepods | 1027 | 894 | AO | TARA_82,83,84,85 |
| 7_20_11 | 180-2000µm | Bilateria | Bilateria | Animals | 1096 | 708 | NAO-SAO | TARA_145,150,151,68,80 |
| 7_3_916 | 180-2000µm | Eumetazoa | Eumetazoa | Animals | 1100 | 379 | MED-NAO | TARA_147,148,149,151,25 |
| 7_4_14 | 180-2000µm | Copepod (Calanoid) | Calanoida | Copepods | 1035 | 294 | MED-NAO | TARA_145,150,152,30 |
| 8_20_12 | 180-2000µm | Copepod (Calanoid) | Calanoida | Copepods | 1164 | 416 | MED-NAO-SAO | TARA_143,144,150,20,22,25,70 |
| 8_50_1 | 180-2000µm | Copepod (Calanoid) | Calanoida | Copepods | 1577 | 639 | MED-NAO-SAO | TARA_145,152,30,66 |
| 8_50_5 | 180-2000µm | Copepod (Calanoid) | Calanoida | Copepods | 1363 | 1237 | MED-NAO | TARA_146,147,148,9 |
| 9_100_5 | 180-2000µm | Bilateria | Bilateria | Animals | 1020 | 667 | NAO | TARA_142,143,144,149,151 |

| MVS name | Size Fraction | Taxonomy (precise) | Retained Taxonomy | Larger group | Number of variants (MVC) | Number of variants (MVS) | Basin(s) | Tara stations |
|---|---|---|---|---|---|---|---|---|
| 10_20_34 | 0.8-5µm | Deltaproteobacteria | Deltaproteobacteria | Bacteria | 1124 | 285 | MED-NAO | TARA_142,147,7,18,22,23 |
| 10_200_9 | 0.8-5µm | Cryptophyta (Geminigeraceae, guillardia theta) | Cryptophyta | Unicellular Eukaryotes | 1121 | 150 | NAO-SAO | TARA_145,146,147,66,67 |
| 10_300_13 | 0.8-5µm | Pelagomonadaceae | Pelagomonadaceae | Unicellular Eukaryotes | 1781 | 1534 | NAO-SAO | TARA_150,152,68,70 |
| 10_300_15 | 0.8-5µm | Alphaproteobacteria (Rhodobactera) | Alphaproteobacteria | Bacteria | 4126 | 1329 | MED | TARA_7,9,18,22,23,25 |
| 12_100_16 | 0.8-5µm | Gammaproteobacteria (Oceanospirillales ) | Gammaproteobacteria | Bacteria | 1444 | 1289 | AO | TARA_82,83,84,85 |
| 12_100_22 | 0.8-5µm | Prochlorococcus (marinus) | Cyanobacteria | Bacteria | 1027 | 297 | MED-NAO-SAO | TARA_150,4,23,25,68,70 |
| 12_200_4 | 0.8-5µm | Alphaproteobacteria | Alphaproteobacteria | Bacteria | 1012 | 268 | MED-NAO-SAO | TARA_142,146,147,18,80 |
| 12_50_5 | 0.8-5µm | Cyanobacteria (Synechococcus) | Cyanobacteria | Bacteria | 1275 | 155 | NAO-SAO | TARA_145,150,151,152,66,68,81 |
| 15_100_9 | 0.8-5µm | Alphaproteobacteria (Candidatus Pelagibacter) | Alphaproteobacteria | Bacteria | 1289 | 511 | MED-NAO-SAO | TARA_142,147,150,151,7,80 |
| 15_300_3 | 0.8-5µm | Virus (Myoviridae) | Virus | Virus | 1085 | 889 | MED-NAO | TARA_150,151,4,18 |
| 15_500_9 | 0.8-5µm | Cyanobacteria (Synechococcus) | Cyanobacteria | Bacteria | 3313 | 898 | MED | TARA_7,9,18,22,23,25 |
| 20_100_3 | 0.8-5µm | Cyanobacteria (Synechococcus) | Cyanobacteria | Bacteria | 1009 | 254 | NAO-SAO | TARA_142,146,147,4,80 |
| 20_100_7 | 0.8-5µm | Cyanobacteria (Synechococcus) | Cyanobacteria | Bacteria | 1022 | 657 | NAO-SAO | TARA_142,146,147,80 |
| 20_200_4 | 0.8-5µm | unclassified | unclassified | Poor classification | 1013 | 322 | MED-NAO-SAO | TARA_4,7,18,23,80 |
| 4_5_2 | 0.8-5µm | Gammaproteobacteria (Cellvibrionales) | Gammaproteobacteria | Bacteria | 1050 | 717 | MED | TARA_9,18,23,25 |
| 4_50_2 | 0.8-5µm | Prymnesiophyceae | Haptophyta | Unicellular Eukaryotes | 2660 | 1204 | AO | TARA_82,83,84,85 |
| 5_100_1 | 0.8-5µm | Micromonas | Mamiellales | Unicellular Eukaryotes | 1089 | 955 | NAO | TARA_142,146,147 |
| 5_20_14 | 0.8-5µm | Gammaproteobacteria | Gammaproteobacteria | Bacteria | 1213 | 103 | MED-NAO | TARA_4,7,9,18,22,23,25 |
| 6_2_5943 | 0.8-5µm | Alphaproteobacteria | Alphaproteobacteria | Bacteria | 1002 | 801 | MED-SAO | TARA_7,9,18,80 |
| 6_5_14 | 0.8-5µm | Bathycoccus | Mamiellales | Unicellular Eukaryotes | 1123 | 353 | NAO-SAO | TARA_145,152,66,67,80,81 |
| 6_9_27 | 0.8-5µm | Alphaproteobacteria | Alphaproteobacteria | Bacteria | 1098 | 404 | AO-MED-SAO | TARA_22,67,80,81,83 |
| 7_100_6 | 0.8-5µm | Flavobacteriia | Flavobacteriia | Bacteria | 1131 | 882 | AO | TARA_82,83,84,85 |
| 7_20_37 | 0.8-5µm | Cyanobacteria (Synechococcus) | Cyanobacteria | Bacteria | 1193 | 917 | MED | TARA_7,9,18,22,23,25 |
| 7_300_4 | 0.8-5µm | Gammaproteobacteria | Gammaproteobacteria | Bacteria | 1061 | 850 | MED | TARA_7,9,18,23,25 |
| 7_7_9 | 0.8-5µm | Cyanobacteria (Synechococcus) | Cyanobacteria | Bacteria | 1005 | 574 | MED-NAO | TARA_142,22,23,25 |
| 8_100_13 | 0.8-5µm | Cyanobacteria (Synechococcus) | Cyanobacteria | Bacteria | 1313 | 1107 | NAO | TARA_146,147,150,151 |
| 8_300_4 | 0.8-5µm | Prymnesiophyceae | Haptophyta | Unicellular Eukaryotes | 1765 | 683 | AO-NAO-SAO | TARA_150,152,68,70,83 |
| 8_300_6 | 0.8-5µm | Alphaproteobacteria (Rhodobacteria) | Alphaproteobacteria | Bacteria | 1496 | 1313 | MED-SAO | TARA_7,9,22,80 |
| 8_5_18 | 0.8-5µm | Cyanobacteria (Synechococcus) | Cyanobacteria | Bacteria | 1007 | 604 | MED-NAO | TARA_145,152,23,81 |
| 8_500_5 | 0.8-5µm | Haptophyta | Haptophyta | Unicellular Eukaryotes | 5629 | 1766 | NAO-SAO | TARA_142,146,147,150,151,68 |
| 8_8_204 | 0.8-5µm | Gammaproteobacteria (Alteromonas) | Gammaproteobacteria | Bacteria | 1002 | 292 | AO-MED-NAO | TARA_4,7,9,82,83 |
| 9_10_185 | 0.8-5µm | Symbiodinium | Dinophyceae | Unicellular Eukaryotes | 1030 | 577 | NAO | TARA_145,146,147,150,151 |
| 9_500_10 | 0.8-5µm | Bathycoccus prasinos | Mamiellales | Unicellular Eukaryotes | 1129 | 712 | NAO-SAO | TARA_146,147,150,80 |
| 10_20_12 | 5-20µm | Copepod (Calanoid?) | Copepodaa | Copepods | 1094 | 174 | MED-NAO-SAO | TARA_150,18,22,25,70,76 |
| 10_200_2 | 5-20µm | Copepod (Calanoid?) | Maxillopoda | Copepods | 1226 | 662 | MED-NAO | TARA_146,147,150,18,25,30 |
| 10_6_36 | 5-20µm | Dinophyceae | Dinophyceae | Unicellular Eukaryotes | 1119 | 656 | NAO-SAO | TARA_150,151,70,76,78 |
| 12_200_5 | 5-20µm | Copepod (Calanoid) | Calanoida | Copepods | 1835 | 1577 | MED-NAO | TARA_147,150,25,30 |
| 12_8_13 | 5-20µm | Copepod (Cyclopoida) | Cyclopoida | Copepods | 1029 | 750 | NAO | TARA_145,147,150,151 |
| 18_200_10 | 5-20µm | Copepod (Calanoid) | Calanoida | Copepods | 1324 | 377 | MED-NAO | TARA_147,149,150,22,25 |
| 18_200_5 | 5-20µm | Copepod (Calanoid) | Calanoida | Copepods | 1073 | 512 | MED-NAO-SAO | TARA_145,152,22,25,66 |
| 18_200_6 | 5-20µm | unclassified | unclassified | Poor classification | 1038 | 489 | NAO | TARA_146,147,148,150,151 |
| 18_300_4 | 5-20µm | Copepod (Oithona) | Cyclopoida | Copepods | 2679 | 1333 | MED-NAO | TARA_151,4,18,25,30 |
| 20_100_6 | 5-20µm | Copepod (Calanoid) | Calanoida | Copepods | 1117 | 616 | MED-NAO | TARA_149,150,151,18,25 |
| 6_6_42 | 5-20µm | Copepod (Calanoid) | Calanoida | Copepods | 1129 | 635 | MED-NAO | TARA_150,4,18,25,30 |
| 7_7_22 | 5-20µm | Dinophyceae | Dinophyceae | Unicellular Eukaryotes | 1091 | 943 | NAO-SAO | TARA_148,70,76,78 |
| 8_10_11 | 5-20µm | Symbiodinium | Dinophyceae | Unicellular Eukaryotes | 1163 | 183 | NAO-SAO | TARA_146,147,148,149,150,70 |
| 8_10_3 | 5-20µm | Haptophyte | Haptophyte | Unicellular Eukaryotes | 1110 | 364 | NAO-SAO | TARA_146,147,148,149,70 |
| 8_7_14 | 5-20µm | Eukaryota | Eukaryota | Poor classification | 1026 | 705 | NAO | TARA_146,147,148,149 |
| 9_100_3 | 5-20µm | Gammaproteobacteria (Alteromonas) | Gammaproteobacteria | Bacteria | 1331 | 218 | MED-NAO-SAO | TARA_4,30,70,76,78 |
| 9_200_1 | 5-20µm | Planctomycetales | Planctomycetales | Bacteria | 1083 | 837 | NAO | TARA_148,150,151,152 |
| 9_4_57 | 5-20µm | Dinophyceae | Dinophyceae | Unicellular Eukaryotes | 1007 | 870 | NAO | TARA_146,147,148,152 |
| 10_100_9 | 20-180µm | Copepod (Oithona) | Cyclopoida | Copepods | 1067 | 124 | MED-NAO-SAO | TARA_144,18,26,30,70,76 |
| 10_6_296 | 20-180µm | Copepod (Calanoid) | Calanoida | Copepods | 1028 | 901 | MED-NAO | TARA_142,143,144,18 |
| 12_100_12 | 20-180µm | Copepod (Cyclopoida) | Cyclopoida | Copepods | 1002 | 552 | NAO-SAO | TARA_72,146,147,148,149,70 |
| 12_3_631 | 20-180µm | Bilateria | Bilateria | Animals | 1027 | 280 | MED-NAO-SAO | TARA_149,150,151,7,78 |
| 12_300_13 | 20-180µm | Copepod (Calanoid) | Calanoida | Copepods | 2084 | 1885 | MED | TARA_11,22,23,26 |
| 12_300_14 | 20-180µm | Copepod | Copepodaa | Copepods | 1219 | 551 | NAO-SAO | TARA_72,142,143,144,76 |
| 15_100_8 | 20-180µm | Maxillopoda | Maxillopoda | Copepods | 1159 | 136 | MED-NAO-SAO | TARA_143,144,150,151,18,22,70,76 |
| 15_200_11 | 20-180µm | Copepod (Calanoid) | Calanoida | Copepods | 1094 | 210 | MED-NAO | TARA_147,149,150,151,152,25,26,30 |

| MVS name | Fixed Effect | Temperature | Salinity | Silicate | Phosphate | Nitrate | Lagrangian | Unexplained | t-SNE_X | t-SNE_Y | cluster |
|---|---|---|---|---|---|---|---|---|---|---|---|
| 15_200_2 | 0.1365 | 0 | 0 | 0 | 0.0021 | 0.0456 | 0 | 0.8156 | 4.47118779172045 | .115172466841 | Unknown |
| 15_300_10 | 7,00E-04 | 0.5499 | 0.0246 | 0.3292 | 0 | 0.0482 | 0.0407 | 0.0068 | 30.1981931594932 | .418526463079 | Temperature |
| 15_500_6 | 0.0909 | 0 | 0 | 0 | 0 | 1,00E-04 | 0 | 0.909 | 4.42664002282318 | .124654105784 | Unknown |
| 15_500_7 | 0.0208 | 0 | 0 | 0.0788 | 0.0357 | 0.0309 | 0.2502 | 0.5836 | 2.55647447734196 | 5.431306888342 | Lagrangian2 |
| 18_300_11 | 0.016 | 6,00E-04 | 0.1951 | 0.0036 | 0.0039 | 0.1102 | 0.335 | 0.3356 | 6.07105491942456 | 12.86877567364 | Lagrangian2 |
| 18_300_9 | 0.0043 | 0 | 0.1084 | 0.0101 | 0.7965 | 0.0381 | 0 | 0.0427 | -24.384163098622 | .130997903207 | Phosphate |
| 20_100_6 | 0.1362 | 3,00E-04 | 2,00E-04 | 0 | 0 | 0 | 0.0427 | 0.8205 | 5.30564992737063 | .339404234582 | Unknown |
| 4_6_23 | 0.0141 | 0.0107 | 0 | 0 | 0 | 0 | 0.8339 | 0.1413 | 3.63067044198188 | 3.59661131617 | Lagrangian |
| 5_10_3 | 0.0283 | 0.0011 | 0.3012 | 0 | 0 | 0.3053 | 0.0816 | 0.2825 | -12.1423379441701 | 24.04254795917 | Nitrate |
| 6_20_9 | 0.0192 | 0 | 0.356 | 0 | 0.1208 | 0.0756 | 0.0257 | 0.4027 | -18.1956956784025 | 20.88128910000 | Salinity |
| 6_50_1 | 0.0964 | 0.0439 | 0.4775 | 0.0706 | 0 | 0 | 0.02 | 0.2916 | -21.271074705172 | 17.25008920196 | Salinity |
| 7_20_9 | 0.0544 | 0.1284 | 0 | 1,00E-04 | 0 | 0 | 0 | 0.817 | 19.0718553049792 | 10.05192023930 | Unknown |
| 7_50_3 | 0.0038 | 0.0085 | 0 | 3,00E-04 | 0.0111 | 0.9533 | 0 | 0.023 | -4.51615987073792 | 29.26216024899 | Nitrate |
| 8_300_3 | 0.0027 | 0 | 0 | 0.1559 | 0.0159 | 0.1615 | 0.6489 | 0.0152 | 1.26922561990608 | 9.63942280294 | Lagrangian |
| 8_50_10 | 0.0297 | 0.0224 | 0 | 0.0307 | 0 | 0.0799 | 0.0047 | 0.8325 | 5.1339622573142 | .643111181006 | Unknown |
| 8_8_91 | 0.0483 | 0.1313 | 0.0723 | 0 | 1,00E-04 | 0 | 0.0223 | 0.7257 | 17.3750702916235 | .899716172106 | Unknown |
| 9_10_18 | 0.0127 | 0.0391 | 0 | 0.0022 | 0.0116 | 0 | 0.7443 | 0.1901 | 3.56427131350743 | 2.27893997009 | Lagrangian |
| 9_100_10 | 0.0333 | 0.0207 | 1,00E-04 | 0 | 2,00E-04 | 6,00E-04 | 0.2448 | 0.7001 | 2.41503697862757 | 4.72242045557 | Lagrangian2 |
| 10_100_10 | 0 | 0 | 0.0042 | 0.5184 | 0.3428 | 0.0956 | 0.039 | 0 | -10.2659206028563 | 3.48043824959 | Silicate |
| 10_20_37 | 0.0844 | 0.0357 | 0.0202 | 0.0106 | 2,00E-04 | 0.0016 | 0 | 0.8473 | 17.1113959103599 | 13.04921214384 | Unknown |
| 10_6_139 | 0.0407 | 0 | 0 | 0.1051 | 3,00E-04 | 0 | 0 | 0.8539 | -4.22662562082439 | 1552145158415 | Silicate |
| 10_8_48 | 0.0235 | 0.0741 | 0.0433 | 1,00E-04 | 0.1471 | 0.0032 | 0.2141 | 0.4945 | 0.99451362229622 | 5.32178912232 | Lagrangian2 |
| 10_9_10 | 0.0095 | 0.0204 | 0.652 | 0.0031 | 1,00E-04 | 1,00E-04 | 0.1719 | 0.143 | -22.9360344918258 | 15.00892516839 | Salinity |
| 10_9_61 | 0.0307 | 0.0015 | 0.0882 | 0 | 0 | 0.0197 | 0 | 0.8599 | 10.1071257364221 | .348474434935 | Unknown |
| 10_9_71 | 0.0579 | 0.0619 | 0.0092 | 2,00E-04 | 0 | 0 | 0 | 0.8708 | 18.0235603825944 | 12.54450738090 | Unknown |
| 12_1_55 | 0.023 | 0.3928 | 0.0666 | 0 | 0.034 | 0 | 1,00E-04 | 0.4834 | 28.0600931751472 | .523705232382 | Temperature |
| 12_10_4 | 0.0296 | 0.0891 | 0.0614 | 1,00E-04 | 0 | 0.4435 | 0.2917 | 0.0847 | -2.0948827310203 | 25.69521221443 | Nitrate |
| 12_100_10 | 0.0364 | 0.0288 | 0.3204 | 0 | 0.0304 | 0.0364 | 0 | 0.5476 | -19.6603042858407 | 21.42445484733 | Salinity |
| 12_100_13 | 0.0199 | 1,00E-04 | 0.3358 | 3,00E-04 | 0 | 0.4442 | 0 | 0.1995 | -11.855478992592 | 24.18232389972 | Nitrate |
| 12_20_30 | 0.053 | 2,00E-04 | 0 | 0.0949 | 0 | 0.0513 | 0.0051 | 0.7955 | -4.42881398539982 | .069023211229 | Silicate |
| 12_200_10 | 0.0212 | 0.3005 | 0.0022 | 0.0796 | 0 | 0 | 0.3838 | 0.2127 | 8.60457351400031 | 9.55110224713 | Lagrangian2 |
| 12_5_104 | 0.0145 | 0.8121 | 0 | 0 | 0.012 | 0 | 0.0158 | 0.1455 | 31.9262270605029 | .234083263668 | Temperature |
| 12_5_123 | 0.0094 | 0.5816 | 0.2451 | 0.0235 | 0 | 0 | 0 | 0.1403 | 31.8791129440348 | .334812060974 | Temperature |
| 12_500_4 | 0.0379 | 1,00E-04 | 0.1318 | 0.2354 | 0.2145 | 0 | 5,00E-04 | 0.3799 | -8.10413179331017 | 2.193175127964 | Silicate |
| 12_500_5 | 0.0457 | 0 | 0 | 6,00E-04 | 0 | 0.0782 | 0.1566 | 0.7188 | 2.40816262308786 | 3.78619691599 | Lagrangian2 |
| 12_8_21 | 0.0019 | 0 | 0 | 0.0378 | 0.6112 | 0.3378 | 0 | 0.0114 | -25.7387195506763 | .810032965142 | Phosphate |
| 15_50_28 | 3,00E-04 | 3,00E-04 | 0.4091 | 0 | 0.4254 | 0.0128 | 0.1502 | 0.0019 | -24.0635929993526 | .7755431158527 | Phosphate |
| 15_9_18 | 0.0831 | 0.0533 | 1,00E-04 | 0 | 0.0299 | 0 | 0 | 0.8335 | 18.6716168324699 | 13.27457276262 | Unknown |
| 15_9_20 | 0.0266 | 1,00E-04 | 0.0853 | 4,00E-04 | 0.3987 | 0.0035 | 0.0877 | 0.3979 | -18.3589807594119 | .414760705473 | Phosphate |
| 18_200_27 | 0.0344 | 0.1316 | 0.0329 | 0 | 0.0511 | 0.0146 | 0.013 | 0.7224 | 18.7838209466465 | 3.355014739256 | Unknown |
| 20_100_10 | 0.0013 | 0.3527 | 0.5096 | 0.0866 | 0 | 0 | 0.0365 | 0.0133 | -19.8624796738073 | 16.74473376787 | Salinity |
| 20_100_9 | 0.0312 | 0.0048 | 1,00E-04 | 0 | 0.0033 | 0 | 0.6483 | 0.3123 | 2.91941969259886 | 1.16438826409 | Lagrangian |
| 20_500_17 | 0 | 0 | 0.7969 | 0 | 0.1512 | 0.0445 | 0.0075 | 0 | -24.7007303315059 | 10.52089764986 | Salinity |
| 20_500_18 | 1,00E-04 | 0.5792 | 0.0773 | 0 | 0 | 0.1041 | 0.2389 | 5,00E-04 | 28.7900893203775 | .782968455677 | Temperature |
| 20_500_21 | 0 | 0.0483 | 0.9063 | 0 | 0.0361 | 0 | 0.0092 | 0 | -25.8565609632823 | 10.94985279534 | Salinity |
| 20_500_22 | 0.0859 | 0 | 0.3974 | 0 | 0 | 9,00E-04 | 0 | 0.5158 | -20.5803474563314 | 20.00098087930 | Salinity |
| 5_20_1 | 0.0401 | 0 | 0 | 0 | 2,00E-04 | 0.3947 | 0.3245 | 0.2405 | -1.18501526633765 | 25.02117485419 | Nitrate |
| 7_20_11 | 0.0211 | 0.5956 | 0.1625 | 0 | 0 | 0 | 0.0087 | 0.2121 | 31.275830834252 | .508564492129 | Temperature |
| 7_3_916 | 0.0624 | 0 | 0.2823 | 2,00E-04 | 7,00E-04 | 0.0204 | 0.0087 | 0.6252 | -19.9787824309686 | 22.44985695548 | Salinity |
| 7_4_14 | 0.1275 | 0 | 0.0448 | 1,00E-04 | 0 | 0.0346 | 0.025 | 0.768 | 10.2641697709783 | .593631282219 | Unknown |
| 8_20_12 | 0.0434 | 0 | 0 | 0.044 | 0 | 0 | 0 | 0.9124 | 1.13416029700442 | .808606920024 | Silicate |
| 8_50_1 | 0.0015 | 3,00E-04 | 0.0782 | 0.7367 | 0.1445 | 0.0298 | 0 | 0.009 | -8.75790224874742 | 0.329637349826 | Silicate |
| 8_50_5 | 0.042 | 0.328 | 3,00E-04 | 0.069 | 0 | 0.3083 | 0 | 0.2524 | 25.1651194819753 | 2.032967102551 | Temperature |
| 9_100_5 | 0.0036 | 0.2402 | 0.0654 | 0.2557 | 0 | 0.1032 | 0.2959 | 0.0359 | 7.36777664234817 | 8.87274596691 | Lagrangian2 |

| MVS name | Fixed Effect | Temperature | Salinity | Silicate | Phosphate | Nitrate | Lagrangian | Unexplained | t-SNE_X | t-SNE_Y | cluster |
|---|---|---|---|---|---|---|---|---|---|---|---|
| 10_20_34 | 0.0433 | 0 | 0.2728 | 0.0127 | 0.0207 | 0 | 0 | 0.6504 | -19.2852283638193 | -23.1444059566222 | Salinity |
| 10_200_9 | 0.0163 | 0.0013 | 0.166 | 4,00E-04 | 0.6519 | 0 | 1,00E-04 | 0.1639 | -24.660485785748 | 4.79460669553445 | Phosphate |
| 10_300_13 | 0.0732 | 0.3063 | 0.0058 | 0 | 0.0818 | 0 | 0.0949 | 0.4379 | 27.3109070506584 | 3.11465514611179 | Temperature |
| 10_300_15 | 0.0612 | 2,00E-04 | 0 | 0 | 0 | 0 | 0.0206 | 0.918 | 4.75275592359791 | 0.377644441444733 | Unknown |
| 12_100_16 | 0.0604 | 0 | 0.5704 | 0.0055 | 2,00E-04 | 5,00E-04 | 8,00E-04 | 0.3622 | -22.5064626462747 | -16.6874182455034 | Salinity |
| 12_100_22 | 0.0599 | 0 | 0 | 0 | 0.0263 | 0.0143 | 0 | 0.8995 | 4.18115102933879 | -3.15751637341172 | Unknown |
| 12_200_4 | 0.004 | 0 | 0.5968 | 8,00E-04 | 0.321 | 0.0118 | 0.0249 | 0.0405 | -24.2004185487957 | -2.13833532078058 | Phosphate |
| 12_50_5 | 0.0343 | 0.0733 | 0.0916 | 0 | 0.003 | 0 | 0.0769 | 0.7209 | 15.5333127120754 | -7.42105856775404 | Unknown |
| 15_100_9 | 0.0147 | 2,00E-04 | 0 | 3,00E-04 | 0.4146 | 0.0975 | 0.2526 | 0.2201 | -18.1895750845467 | 4.72573310338044 | Phosphate |
| 15_300_3 | 0.0067 | 9,00E-04 | 0.0013 | 0.2456 | 0.0131 | 0.3123 | 0.3799 | 0.0404 | 0.239260309422452 | -24.0797471356567 | Nitrate |
| 15_500_9 | 0.0179 | 0 | 2,00E-04 | 0 | 0.0037 | 0.448 | 0.2616 | 0.2686 | -1.12923237020809 | -26.0411490685487 | Nitrate |
| 20_100_3 | 0.0104 | 0.4698 | 0 | 6,00E-04 | 0.0021 | 0.4125 | 0 | 0.1045 | 25.3604386974162 | -1.83895881863454 | Temperature |
| 20_100_7 | 0.0089 | 0.9136 | 1,00E-04 | 0.0068 | 0.0055 | 0.0089 | 0.0019 | 0.0543 | 31.9238061463571 | 10.3408775262454 | Temperature |
| 20_200_4 | 0.0428 | 0 | 0 | 0.0172 | 1,00E-04 | 0.0536 | 0.4573 | 0.429 | 3.51240807578828 | 20.4753395787973 | Lagrangian2 |
| 4_5_2 | 0 | 0.0058 | 0.0033 | 0.7431 | 0.0075 | 0.0227 | 0.2176 | 0 | -7.7425785938013 | 11.4199653773324 | Silicate |
| 4_50_2 | 0 | 0.0095 | 0.0195 | 0.0081 | 0.2029 | 0 | 0.76 | 0 | 1.12165258773066 | 32.5442334282814 | Lagrangian |
| 5_100_1 | 0 | 0 | 0 | 0 | 2,00E-04 | 0.9998 | 0 | 0 | -4.82587939887527 | -29.1194096160284 | Nitrate |
| 5_20_14 | 0.021 | 0 | 0.0032 | 0.0069 | 0.0231 | 0.0552 | 0.4496 | 0.441 | 3.09189733958743 | 20.2691689436746 | Lagrangian2 |
| 6_2_5943 | 0.0014 | 0 | 0 | 9,00E-04 | 0 | 0.0371 | 0.9519 | 0.0086 | 3.49402719457801 | 35.0509925262005 | Lagrangian |
| 6_5_14 | 0.0488 | 0.0582 | 1,00E-04 | 0.1553 | 1,00E-04 | 6,00E-04 | 0.0026 | 0.7343 | -5.56638269710948 | 0.622409323502679 | Silicate |
| 6_9_27 | 0.0083 | 0.1675 | 0 | 0.0718 | 0.6538 | 0.0017 | 0.0141 | 0.0827 | -22.8530316585461 | 5.90208665408954 | Phosphate |
| 7_100_6 | 0.1424 | 0 | 5,00E-04 | 0 | 0 | 0.0024 | 0 | 0.8547 | 3.83901103608941 | -1.7493423067805 | Unknown |
| 7_20_37 | 0.0561 | 0.0061 | 0.0629 | 1,00E-04 | 2,00E-04 | 3,00E-04 | 0.0324 | 0.8421 | 11.0312126026875 | -8.18142113801941 | Unknown |
| 7_300_4 | 0.047 | 0.0212 | 0.1711 | 1,00E-04 | 0 | 0.0066 | 0.2823 | 0.4718 | 5.68920529688963 | 13.1286904871818 | Lagrangian2 |
| 7_7_9 | 0.0044 | 0 | 0.4754 | 0 | 0 | 3,00E-04 | 0.4933 | 0.0266 | 7.21337188302061 | 11.7082587625771 | Lagrangian2 |
| 8_100_13 | 1,00E-04 | 0 | 0 | 0.8752 | 0.1232 | 0.001 | 0 | 6,00E-04 | -8.06111905726165 | 10.2568634550825 | Silicate |
| 8_300_4 | 5,00E-04 | 0 | 0.8883 | 0.0721 | 0.0346 | 0 | 0 | 0.0046 | -25.0095047157473 | -11.5838267001138 | Salinity |
| 8_300_6 | 0.0097 | 0 | 0 | 1,00E-04 | 0 | 0 | 0.9323 | 0.058 | 4.16244004821243 | 34.6775330526547 | Lagrangian |
| 8_5_18 | 0.1192 | 2,00E-04 | 1,00E-04 | 0.1243 | 3,00E-04 | 0.0412 | 0 | 0.7148 | -4.90096789392655 | -0.516402461301072 | Silicate |
| 8_500_5 | 0.0084 | 0.6347 | 0 | 0 | 0.1825 | 0 | 0.0486 | 0.1258 | 30.3251818866403 | 7.67820944013163 | Temperature |
| 8_8_204 | 0.0448 | 0 | 0 | 0.112 | 0 | 5,00E-04 | 0.3957 | 0.4471 | 4.03896719108721 | 19.0029510970795 | Lagrangian2 |
| 9_10_185 | 0.0909 | 0 | 0 | 0 | 0 | 0 | 0 | 0.909 | 3.03084898465676 | -1.36339573546197 | Unknown |
| 9_500_10 | 0.0015 | 0 | 0 | 0.5148 | 0 | 0 | 0.4747 | 0.009 | -6.93128868749637 | 12.4976975191702 | Silicate |
| 10_20_12 | 0.0578 | 0.0024 | 0.0057 | 1,00E-04 | 0 | 0.0656 | 1,00E-04 | 0.8684 | 4.65512275720554 | -6.67291439744875 | Unknown |
| 10_200_2 | 0.0429 | 0 | 0.0384 | 0.2471 | 0.0268 | 0 | 0 | 0.6447 | -6.83326226771352 | 1.34517529420826 | Silicate |
| 10_6_36 | 0.0213 | 0.1147 | 0 | 0 | 0 | 0 | 0.6506 | 0.2134 | 3.83370080819903 | 30.9339557153637 | Lagrangian |
| 12_200_5 | 0.0057 | 0.1381 | 0 | 0 | 0 | 0.5755 | 0.2469 | 0.0339 | -2.38461405586291 | -26.8214199114312 | Nitrate |
| 12_8_13 | 0.0087 | 0.0035 | 0.9332 | 0.0024 | 0 | 0 | 0 | 0.0522 | -25.940171003814 | -11.87760360972 | Salinity |
| 18_200_10 | 0.0819 | 0.087 | 5,00E-04 | 3,00E-04 | 0 | 0.007 | 4,00E-04 | 0.8228 | 18.7160988608554 | -11.5249284390853 | Unknown |
| 18_200_5 | 4,00E-04 | 0 | 2,00E-04 | 0.3852 | 0.5143 | 0 | 0.0956 | 0.0042 | -10.7792925801925 | 8.04565681269603 | Silicate |
| 18_200_6 | 0.066 | 1,00E-04 | 0.2429 | 0 | 0.0242 | 0 | 0.0063 | 0.6605 | -19.9143614476248 | -23.9787545407648 | Salinity |
| 18_300_4 | 0.0363 | 0.1037 | 0 | 0.0953 | 0.0206 | 2,00E-04 | 0.378 | 0.3658 | 4.9685251698207 | 18.9167861876584 | Lagrangian2 |
| 20_100_6 | 0.0216 | 0.0124 | 0.3192 | 0 | 0.4044 | 0.0022 | 0.0247 | 0.2155 | -23.970068629715 | -0.0398704597345631 | Phosphate |
| 6_6_42 | 0.0322 | 0 | 4,00E-04 | 0.0991 | 0 | 0.3801 | 0.1662 | 0.3219 | -0.100085196037452 | -25.7116746995848 | Nitrate |
| 7_7_22 | 0.022 | 0.7535 | 0.0618 | 0 | 0.0307 | 0 | 0 | 0.132 | 31.6791186593824 | 8.49484111212348 | Temperature |
| 8_10_11 | 0.009 | 0 | 0.1604 | 0 | 0.6959 | 0 | 0 | 0.1346 | -23.9773677901634 | 5.0722341310757 | Phosphate |
| 8_10_3 | 0.036 | 0.5173 | 1,00E-04 | 0 | 1,00E-04 | 1,00E-04 | 0.0857 | 0.3607 | 29.1142194118142 | 4.63904410710441 | Temperature |
| 8_7_14 | 0.0183 | 0.4254 | 0.0021 | 4,00E-04 | 0 | 0 | 0.4438 | 0.1099 | 9.291340475905 | 19.8150356098709 | Lagrangian2 |
| 9_100_3 | 0.0319 | 0.0131 | 0.5573 | 0.0026 | 0 | 0 | 0.0762 | 0.3189 | -22.3121267676777 | -15.9593884437398 | Salinity |
| 9_200_1 | 0.0311 | 4,00E-04 | 2,00E-04 | 0.0013 | 0.0765 | 0.0475 | 0.6583 | 0.1848 | 2.05188680125501 | 31.2465942500519 | Lagrangian |
| 9_4_57 | 0.0114 | 0.9202 | 0 | 1,00E-04 | 0 | 0 | 0 | 0.0683 | 32.8172520078053 | 9.95399878971633 | Temperature |
| 10_100_9 | 0.0408 | 1,00E-04 | 0.1122 | 1,00E-04 | 0.2331 | 0 | 0 | 0.6138 | -17.5888110134198 | 2.47530094125205 | Phosphate |
| 10_6_296 | 0.1159 | 0.1415 | 0 | 3,00E-04 | 0.0466 | 0 | 0.0017 | 0.694 | 19.5172553420878 | -8.88862583022718 | Unknown |
| 12_100_12 | 0.0146 | 0 | 0.187 | 6,00E-04 | 0.558 | 0.0211 | 0 | 0.2187 | -24.5569796869584 | 3.87690230164111 | Phosphate |
| 12_3_631 | 0.0643 | 0 | 0.2843 | 0.0084 | 0 | 0 | 0 | 0.6429 | -20.6990742399602 | -22.9329484421086 | Salinity |
| 12_300_13 | 0.0859 | 0 | 0 | 0 | 0.3986 | 0 | 2,00E-04 | 0.5153 | -18.80780024908 | 3.03281612988102 | Phosphate |
| 12_300_14 | 0.0838 | 0 | 0.0652 | 0.0102 | 0 | 0 | 0.0012 | 0.8396 | 10.9302001997217 | -9.11432651261761 | Unknown |
| 15_100_8 | 0.0212 | 0.0807 | 0.1232 | 9,00E-04 | 3,00E-04 | 0.1519 | 0.0275 | 0.5944 | 6.98907704822802 | -9.07909278962472 | Unknown |
| 15_200_11 | 0.0345 | 0 | 1,00E-04 | 0 | 0 | 0 | 0 | 0.9654 | 3.49919108024647 | -0.581716262594579 | Unknown |

| MVS name | Global Median-FST | Maximum-pairwise-FST | Minimum-pairwise-FST |
|---|---|---|---|
| 15_200_2 | 1 | 1 | 0.0668117397611873 |
| 15_300_10 | 0.0663472727891968 | 0.065474365525465 | 0.019770203930815 |
| 15_500_6 | 0.521612311048298 | 1 | 0.105263157894737 |
| 15_500_7 | 0.086066097730318 | 0.0869565217391303 | 0.0238095238095236 |
| 18_300_11 | 0.0792052880887083 | 0.0875011984276626 | 0.0182185932150796 |
| 18_300_9 | 0.0734058728819302 | 0.0779220779220779 | 0.0200421455586124 |
| 20_100_6 | 0.067463727587513 | 0.0822539699861012 | 0.0136054421768702 |
| 4_6_23 | 0.155837686929212 | 0.2 | 0.0409386925732195 |
| 5_10_3 | 0.147723180945268 | 0.211111111111111 | 0.0952380952380952 |
| 6_20_9 | 0.112797768599575 | 0.119429590017825 | 0.0412355090738916 |
| 6_50_1 | 1 | 1 | 0.04423109822983 |
| 7_20_9 | 0.62203461845416 | 1 | 0.0586300192363809 |
| 7_50_3 | 0.829383319847837 | 1 | 0.167224080267559 |
| 8_300_3 | 0.132090177321135 | 0.125874125874126 | 0.03125 |
| 8_50_10 | 0.115153358074095 | 0.111111111111111 | 0.0327472227901587 |
| 8_8_91 | 0.192030964412429 | 0.17979797979798 | 0.0552637588490521 |
| 9_10_18 | 0.148449374569555 | 0.156739811912226 | 0.054054054054054 |
| 9_100_10 | 0.148590000251264 | 0.156299840510367 | 0.0484848484848486 |
| 10_100_10 | 0.0484524164559834 | 0.0560437738351849 | 0.00844805431198999 |
| 10_20_37 | 0.0649821379555036 | 0.0651890482398958 | 0.0194959644500929 |
| 10_6_139 | 0.0969840114646646 | 0.0740740740740736 | 0.0346543756953051 |
| 10_8_48 | 0.0960203508981884 | 0.13620495430293 | 0.0207893622201339 |
| 10_9_10 | 0.136978428428559 | 0.163791985190169 | 0.0444409289653504 |
| 10_9_61 | 0.0963948406746777 | 0.0830564784053157 | 0.0229310180376512 |
| 10_9_71 | 0.0822601249654252 | 0.0745156482861403 | 0.0264603727491431 |
| 12_1_55 | 0.0672242899570278 | 0.0603150351049511 | 0.0217391304347826 |
| 12_10_4 | 0.0313788826634694 | 0.0284288608805309 | 0.0170940170940171 |
| 12_100_10 | 0.128756822460663 | 0.129845766028724 | 0.0470633803967136 |
| 12_100_13 | 0.109147060686705 | 0.121390656408882 | 0.0247209171474904 |
| 12_20_30 | 0.0894816130001802 | 0.11010101010101 | 0.021914489401471 |
| 12_200_10 | 0.0790602454942032 | 0.0737327188940094 | 0.0304652689026287 |
| 12_5_104 | 0.0613485409739657 | 0.0834042553191486 | 0.0211290362958716 |
| 12_5_123 | 0.0673241286299597 | 0.0625000000000002 | 0.0173913043478259 |
| 12_500_4 | 0.0789890908152684 | 0.0625000000000002 | 0.0241686001546791 |
| 12_500_5 | 0.0716924186526509 | 0.0768049155145926 | 0.01556379981276 |
| 12_8_21 | 0.0555360737963558 | 0.0440770749156407 | 0.030811990210267 |
| 15_50_28 | 0.0448159591916502 | 0.0430347897472669 | 0.0189385416502271 |
| 15_9_18 | 0.086624779012917 | 0.135309791430584 | 0.0141403405294573 |
| 15_9_20 | 0.0709683489850122 | 0.0715276401822873 | 0.0241189421300933 |
| 18_200_27 | 0.0768260616630634 | 0.0820081781008633 | 0.0196644598654653 |
| 20_100_10 | 0.105175875170361 | 0.146479570099366 | 0.0220450137814909 |
| 20_100_9 | 0.0868416099616802 | 0.0914973363947405 | 0.0261319299519103 |
| 20_500_17 | 0.17377650462589 | 0.303030303030303 | 0.0426693975081074 |
| 20_500_18 | 0.187957588970585 | 0.230769230769231 | 0.0647773279352225 |
| 20_500_21 | 0.0400288692872349 | 0.0402061022428771 | 0.0120834609735718 |
| 20_500_22 | 0.0473639240018329 | 0.0518484430127322 | 0.0152195451000234 |
| 5_20_1 | 0.349880434359205 | 0.608724104205566 | 0.0444444444444442 |
| 7_20_11 | 0.0810433281494406 | 0.0985424655501845 | 0.0167627029858563 |
| 7_3_916 | 0.0626440435192702 | 0.0855614973262031 | 0.0215478870878488 |
| 7_4_14 | 0.292682926829269 | 0.384615384615384 | 0.128941499399554 |
| 8_20_12 | 0.0968668542577331 | 0.0876949071363954 | 0.0186055407486239 |
| 8_50_1 | 1 | 1 | 0.0976190476190476 |
| 8_50_5 | 0.0490385160866168 | 0.0381157540720548 | 0.0267034962902843 |
| 9_100_5 | 0.0730653071972875 | 0.0771567735111065 | 0.0182329846290703 |

| MVS name | Global Median-FST | Maximum-pairwise-FST | Minimum-pairwise-FST |
|---|---|---|---|
| 10_20_34 | 0.212172308965259 | 0.287390029325514 | 0.065952792979559 |
| 10_200_9 | 0.586440195135847 | 1 | 0.101738410596027 |
| 10_300_13 | 0.200447315740737 | 0.181244475294075 | 0.106186924089513 |
| 10_300_15 | 0.148699815326744 | 0.222222222222222 | 0.0490795874795046 |
| 12_100_16 | 0.356020942408377 | 0.50830632859517 | 0.026431718061674 |
| 12_100_22 | 0.120025630019471 | 0.145583902714223 | 0.0277010736464136 |
| 12_200_4 | 0.239471144505039 | 0.537884400954263 | 0.0450006656903206 |
| 12_50_5 | 0.258187761895973 | 0.507142857142857 | 0.0779220779220779 |
| 15_100_9 | 0.109202798046532 | 0.0952380952380954 | 0.0475654376583173 |
| 15_300_3 | 0.314050833685342 | 0.484622553588071 | 0.0431355287903641 |
| 15_500_9 | 0.0892665703349781 | 0.0983785256846985 | 0.0275385492044306 |
| 20_100_3 | 0.541967854122916 | 1 | 0.0408163265306123 |
| 20_100_7 | 0.0967741935483874 | 0.139330723135315 | 0.0242389233438191 |
| 20_200_4 | 0.107195706387299 | 0.133333333333333 | 0.0291278138535278 |
| 4_5_2 | 1 | 1 | 0.08 |
| 4_50_2 | 0.840165150225006 | 1 | 0.0488745657343849 |
| 5_100_1 | 0.22657229710654 | 0.351465528143954 | 0.0728744939271255 |
| 5_20_14 | 0.366731173546076 | 0.940350877192982 | 0.0606060606060606 |
| 6_2_5943 | 0.16984984984985 | 0.271939736346515 | 0.0414250207125106 |
| 6_5_14 | 0.41132616068653 | 0.666666666666667 | 0.11017661017661 |
| 6_9_27 | 0.379888887671156 | 0.666666666666667 | 0.0726733407489804 |
| 7_100_6 | 0.582984106831389 | 0.81917211328976 | 0.0869565217391301 |
| 7_20_37 | 0.0989243401614111 | 0.115299141574797 | 0.028808397541274 |
| 7_300_4 | 0.232256694075769 | 0.222222222222222 | 0.0555395201026268 |
| 7_7_9 | 0.147290081390764 | 0.329113924050633 | 0.0139612717519871 |
| 8_100_13 | 0.154651651645635 | 0.204081632653061 | 0.037648203253299 |
| 8_300_4 | 1 | 1 | 0.0869565217391304 |
| 8_300_6 | 0.792941306392744 | 1 | 0.074074074074074 |
| 8_5_18 | 0.395740679877601 | 0.76923076923077 | 0.118518518518519 |
| 8_500_5 | 0.1794950829756 | 0.22637453493179 | 0.0444444444444442 |
| 8_8_204 | 0.126734547327716 | 0.117528405604062 | 0.019953547297297 |
| 9_10_185 | 0.0593095994725671 | 0.0645161290322581 | 0.0127323656735423 |
| 9_500_10 | 0.155517847040633 | 0.235978733095142 | 0.0234193178910428 |
| 10_20_12 | 0.0830365518266337 | 0.0831598468533141 | 0.0246059974251589 |
| 10_200_2 | 0.104337169061364 | 0.0824866514906724 | 0.0417198808003404 |
| 10_6_36 | 0.192948361028995 | 0.281505659824661 | 0.0385494954276136 |
| 12_200_5 | 0.082963760117591 | 0.0571428571428572 | 0.0406795655102379 |
| 12_8_13 | 0.218905472636816 | 0.303030303030303 | 0.0564516129032257 |
| 18_200_10 | 0.0843962978332418 | 0.125 | 0.0189133232611494 |
| 18_200_5 | 0.206469295782867 | 0.405670772956589 | 0.0322945108746822 |
| 18_200_6 | 0.0645701623559973 | 0.0744936280243803 | 0.018104454082591 |
| 18_300_4 | 0.0650893472190191 | 0.0698918875333686 | 0.0244565217391304 |
| 20_100_6 | 0.0706692956112596 | 0.0559350459782924 | 0.028509181697905 |
| 6_6_42 | 0.0842365423255101 | 0.105263157894737 | 0.032247701933225 |
| 7_7_22 | 0.0707184157528251 | 0.0622868114640334 | 0.0322997416020671 |
| 8_10_11 | 0.0996631783175812 | 0.112869771991629 | 0.0192307692307695 |
| 8_10_3 | 0.230730024500979 | 0.307828865613349 | 0.0373370206243403 |
| 8_7_14 | 0.067441807895049 | 0.0588235294117644 | 0.0260133309313637 |
| 9_100_3 | 0.259117609455935 | 0.278129395218002 | 0.117647058823529 |
| 9_200_1 | 0.319925493248406 | 0.571428571428571 | 0.0551750380517502 |
| 9_4_57 | 0.136765120012588 | 0.180759046778464 | 0.0350877192982459 |
| 10_100_9 | 0.591419707826614 | 1 | 0.072463768115942 |
| 10_6_296 | 0.0652864419283524 | 0.0444444444444445 | 0.0325312542008335 |
| 12_100_12 | 0.0844644521957899 | 0.08 | 0.033045787609369 |
| 12_3_631 | 0.0833242428899492 | 0.081447963800905 | 0.0318138614172483 |
| 12_300_13 | 0.064807474366638 | 0.0651890482398958 | 0.0253069828721996 |
| 12_300_14 | 0.0639722985402082 | 0.051992726915327 | 0.0282718437912304 |
| 15_100_8 | 0.082596343254904 | 0.0995581737849778 | 0.0255254571667044 |
| 15_200_11 | 0.0974358872892014 | 0.0833333333333333 | 0.0292712626451852 |
